# Ultrasound Transparent Neural Interfaces for Multimodal Interaction

**DOI:** 10.1101/2025.07.14.664713

**Authors:** Raphael Panskus, Andrada I. Velea, Lukas Holzapfel, Christos Pavlou, Qingying Li, Chaoyi Qin, Flora M. Nelissen, Rick Waasdorp, David Maresca, Valeria Gazzola, Vasiliki Giagka

## Abstract

Neural interfaces that unify diagnostic and therapeutic functionalities hold particular promise for advancing both fundamental neuroscience and clinical neurotechnology. Functional ultrasound imaging (fUSI) has recently emerged as a powerful modality for high-resolution, non-invasive monitoring of brain function and structure. However, conventional metal-based microelectrodes typically impede ultrasound propagation, limiting compatibility with fUSI. Here, we present flexible, ultrasound-transparent neural interfaces that retain practical metal thicknesses while achieving high acoustic transparency. We introduce a theoretical and simulation-based framework to investigate the conditions under which commonly used polymers and metals in neural interfaces can become acoustically transparent. Based on these insights, we propose design guidelines that maximize ultrasound transmission through soft neural interfaces. We experimentally validate our approach through immersion experiments and by demonstrating the acoustic transparency of a suitably engineered interface using fUSI in phantom and in vivo experiments. Finally, we discuss the potential extension of this approach to therapeutic focused ultrasound (FUS). This work establishes a foundation for the development of multimodal neural interfaces with enhanced diagnostic and therapeutic capabilities, enabling both scientific discovery and translational impact.

## Introduction

Neural interfaces are essential tools in neuroscience and medicine, enabling insights into brain function and new therapies for neurological disorders^1,2^. Electrophysiology, long considered the gold standard for studying neuronal activity, offers direct access to the brain dynamics but remains limited in spatial coverage^3^. Both rigid and flexible electrodes struggle to capture large-scale activity across distributed neural networks. To address this, researchers have begun integrating complementary, modalities^4–8^ such as functional magnetic resonance imaging (fMRI)^9^, two-photon imaging^10^ or optogenetics^11^, made possible by magnetically- and optically-compatible or transparent neuroelectronic implants^12–15^. However, optical methods face translational barriers due to limited penetration and the need for genetic modification^16^. fMRI systems are constrained by size, high cost, and susceptibility to metal-induced image artefacts, which hinder integration with implanted electrodes and reduce portability. Meanwhile, ultrasound has re-emerged as a powerful, non-invasive tool for brain imaging and neuromodulation^17–20^. Innovations like ultrafast ultrasound^21^ and novel contrast agents^22^ have enabled functional ultrasound imaging (fUSI), which maps hemodynamic changes correlated with neural activity, thus offering an indirect but real-time visualization of brain activity. fUSI provides a uniquely advantageous combination of wide spatial coverage, high spatiotemporal resolution, and high sensitivity, offering a closer link to underlying neuronal population dynamics than fMRI^23^ while remaining minimally invasive and portable^24^. When fUSI is combined with electrophysiology, it allows researchers to correlate direct neural signals with hemodynamic responses, enabling a more comprehensive understanding of brain activity^25,26^. Additionally, the potential of ultrasound for targeted neuromodulation opens new therapeutic opportunities. Integrating electrophysiology helps clarify how specific ultrasound parameters affect neural circuits, addressing still-unclear mechanisms^27–30^.

To fully realise the potential of these multimodal approaches, however, neural interfaces must be compatible with ultrasound. Conventional approaches based on rigid substrates or thicker metal-based electrodes often obstruct acoustic waves, limiting the integration of electrophysiology with functional or focused ultrasound. This creates a need for neural interfaces that are both flexible and acoustically transparent, enabling seamless combination of electrical recording with ultrasound-based imaging or neuromodulation (Fig. 1). While certain flexible metal-based neural electrodes can achieve partial ultrasound compatibility, the conditions under which traditional neural interfaces, particularly those using metal-based conductors, can be made ultrasound-transparent remain largely unexplored. As we demonstrate later in this work, appropriately engineered electrode arrays can meet these conditions. Recently, graphene micro-transistor arrays have demonstrated compatibility with fUSI^31^. The atomic-scale thickness of graphene inherently provides it with exceptional acoustic transparency, making it an attractive material for integration with ultrasound-based modalities. However, in most practical implementations, only the electrode contacts are composed of graphene, while the interconnects and traces remain metal-based, and can often still impede ultrasound propagation. Other studies^32^ have examined polymers to create acoustically transparent windows.

**Figure 1.**
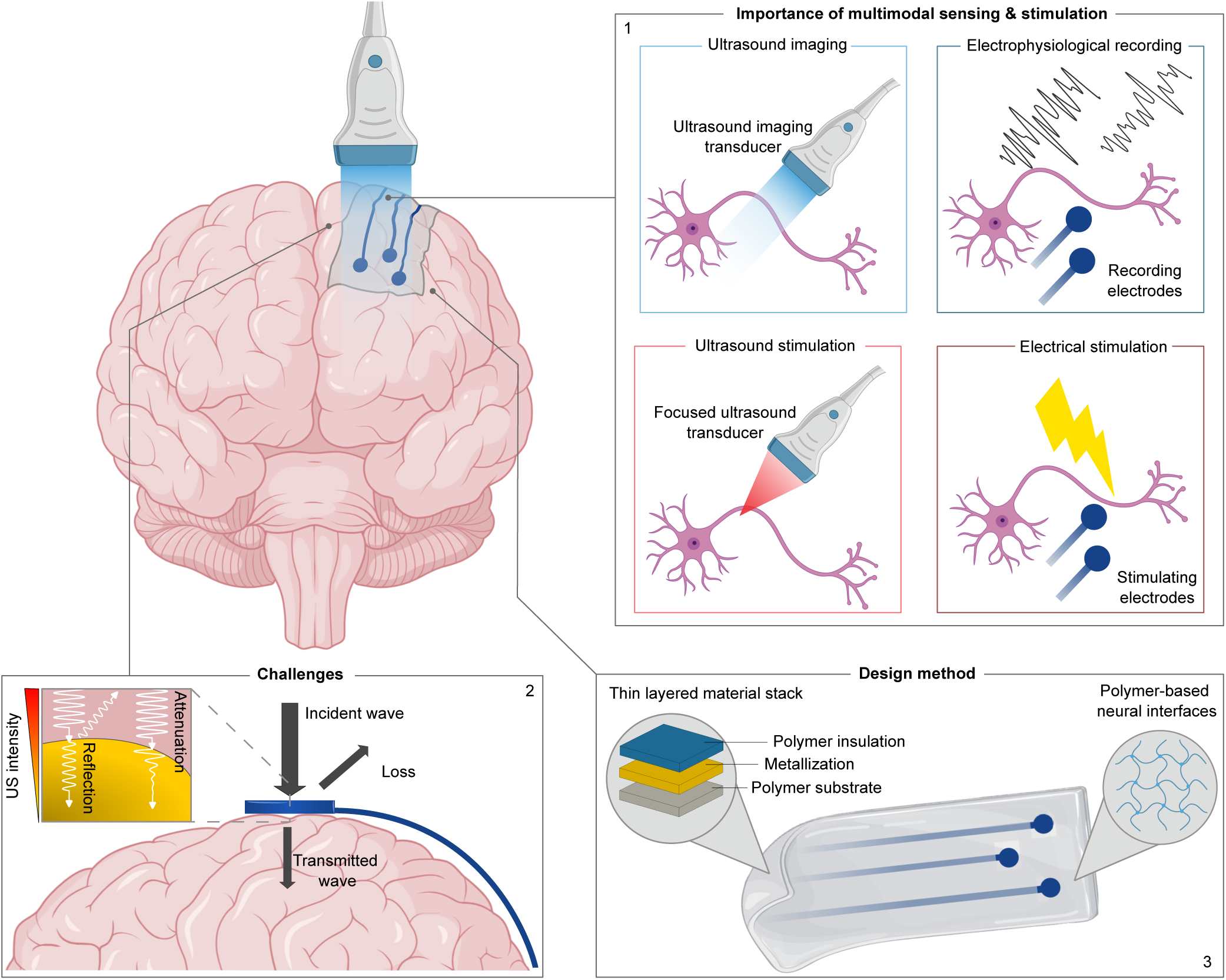
Schematic illustration of the concept of ultrasound-compatible neural electrodes. (1) Importance of multimodal sensing and stimulation, (2) Challenges of ultrasound wave propagation, (3) Design method for ultrasound transparent neural electrodes (inspired by^72^)^73^.

Yet, the conditions under which such multilayer structures, incorporating metals and polymers with varying properties, achieve effective transparency have not been systematically investigated. As a result, the acoustic performance of complete flexible, metal-based neural interfaces remains poorly understood. In this study we address this gap by systematically investigating and validating design strategies for enhancing ultrasound transmission across a complete neural interface. Rather than proposing a single device architecture, we introduce a modelling and validation framework that can be broadly applied to optimise compatibility with ultrasound imaging modalities across a wide class of flexible neural interfaces (Fig. 1).

Using a numerical model, we systematically investigate how medical-grade polymers and commonly used metals in neural implants can be configured to achieve acoustic transparency. We validate our modelling approach through controlled immersion experiments with layered materials representing neural interface stacks. Finally, we demonstrate that suitably engineered polymer–metal-based neural interfaces retain compatibility with fUSI, including in vivo functional ultrasound Doppler imaging of the mouse brain through the implanted interface.

## Results

### Numerical simulations

Ultrasound transmission through layered materials is influenced by acoustic impedance mismatches at interfaces and energy loss due to absorption within the materials. To quantify these effects, we employed the transfer matrix method (TMM), a standard analytical technique for modeling wave propagation in stratified media^33,34^. In our model, we consider a stack of *n* distinct and homogeneous layers positioned between two semi-infinite media, with water used as a proxy for tissue (Fig. 3a). TMM relates the acoustic pressure and particle velocity at the input and output of the stack, allowing us to compute the transmission coefficient across the structure. The transmission coefficient represents the ratio of transmitted to incident pressure amplitudes, reflecting how efficiently an acoustic wave passes through the entire stack. In this work, we report the magnitude of the transmission coefficient |*T*|, which quantifies the proportion of acoustic pressure that is transmitted.

The transmission depends on both the intrinsic wave properties—such as frequency *f* (or angular frequency *ω*) and wavenumber *k_i_*, and the physical characteristics of each layer, including longitudinal wave velocity *c_i_*, density *ρ_i_*, thickness *t_i_* and attenuation coefficient *α_i_*. Interfacial behaviour is governed by acoustic impedance mismatches, where the acoustic impedance of each layer is given by *Z_i_* = *ρ_i_c_i_*.

Flexible neural interfaces typically consist of a polymer substrate, a metallic conductor, and an insulating layer. Based on the material properties, we evaluated the relative advantages and limitations of each material against three key criteria that typically guide the design of ultrasound-enabled bioelectronic implants: acoustic impedance, acoustic attenuation, and Young’s modulus (Fig. 2). Fig. 2b details the specific material parameters used in this study, including water, Platilon 4201AU thermoplastic polyurethane (TPU), MED2-4213 medical-grade polydimethylsiloxane (PDMS), LTC9320 polyimide (PI), parylene-C (ParC), titanium (Ti), platinum (Pt), and gold (Au). Ideally, ultrasound-compatible neural electrodes should maximize acoustic transmission while maintaining sufficient mechanical stability and low flexural rigidity. Notably, the thickness of the metallization layer can further influence the electrical performance and must therefore be considered during design optimization. For numerical modeling, electrode regions with exposed sites are approximated as a three-layer stack comprising substrate, metallic adhesion layer, and electrode contact. Embedded tracks were modeled as a four-layer structure with an additional insulation layer. Polymer layers were treated as acoustically lossy and assigned frequency-dependent attenuation values, while metal layers were considered acoustically lossless. This approach allowed us to capture the primary effects influencing ultrasound transmission through soft, polymer–metal multilayer stacks representative of neural interfaces. Further methodological details are provided in the Methods section.

**Figure 2.**
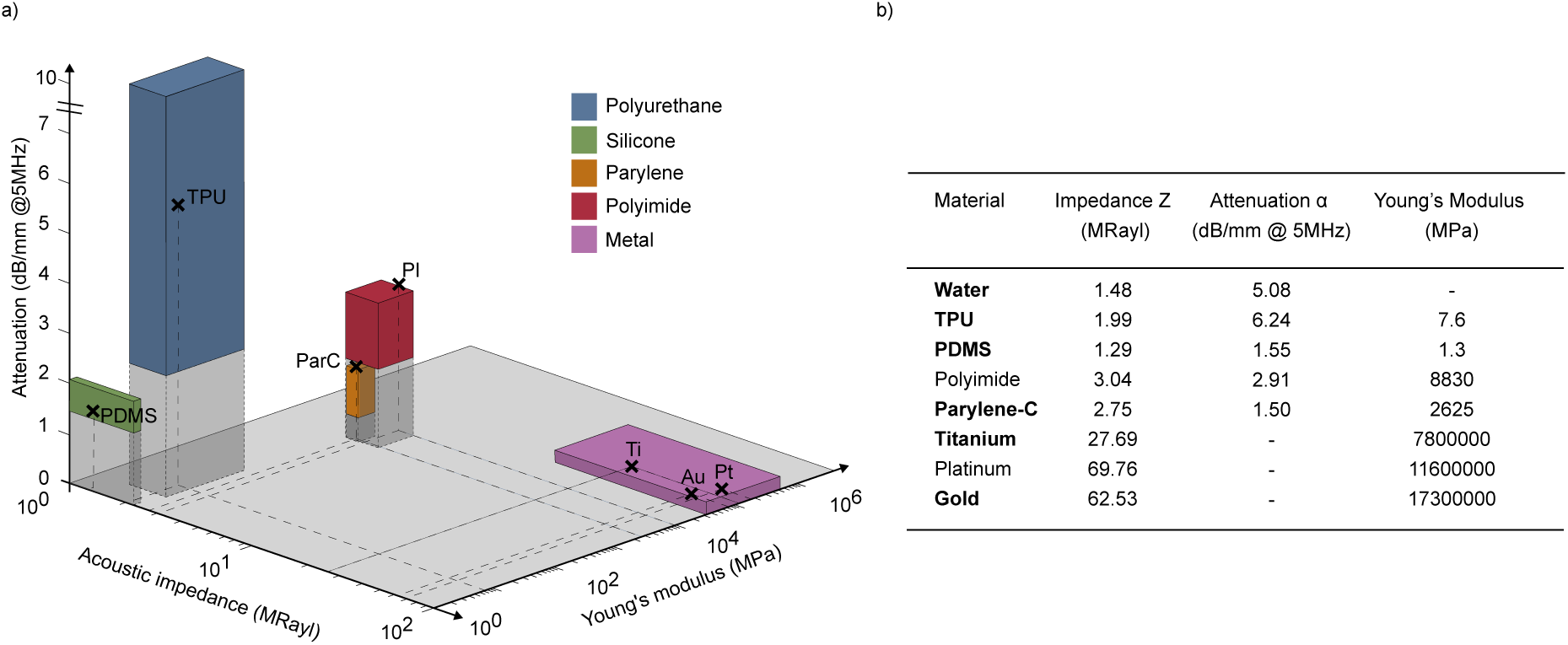
Comparison of material properties relevant for ultrasound-compatible neural interfaces. a) Acoustic impedance, acoustic attenuation (at 5 MHz), and Young’s modulus are shown for representative materials commonly used for neural interfaces. Materials include polymers and metals. Shaded volumes indicate approximate ranges of reported values, while markers represent representative data points obtained from literature^74–99^. Polymers generally exhibit lower acoustic impedance, resulting in reduced ultrasound reflection at tissue interfaces, but tend to have higher acoustic attenuation, leading to greater energy losses. Metals, in contrast, provide superior mechanical robustness but poor ultrasound transmission due to high impedance mismatch. Overall, implant design requires balancing material thickness and selection based on acoustic impedance (to minimise reflections), attenuation (to reduce energy losses), and Young’s modulus (to ensure sufficient mechanical stability and low flexural rigidity). b) Relevant simulation parameters of the selected materials. Materials highlighted in bold represent the specific parameter values selected for the simulations conducted in this study. Additional material simulations can be found in Supplementary Fig. 1.

**Figure 3.**
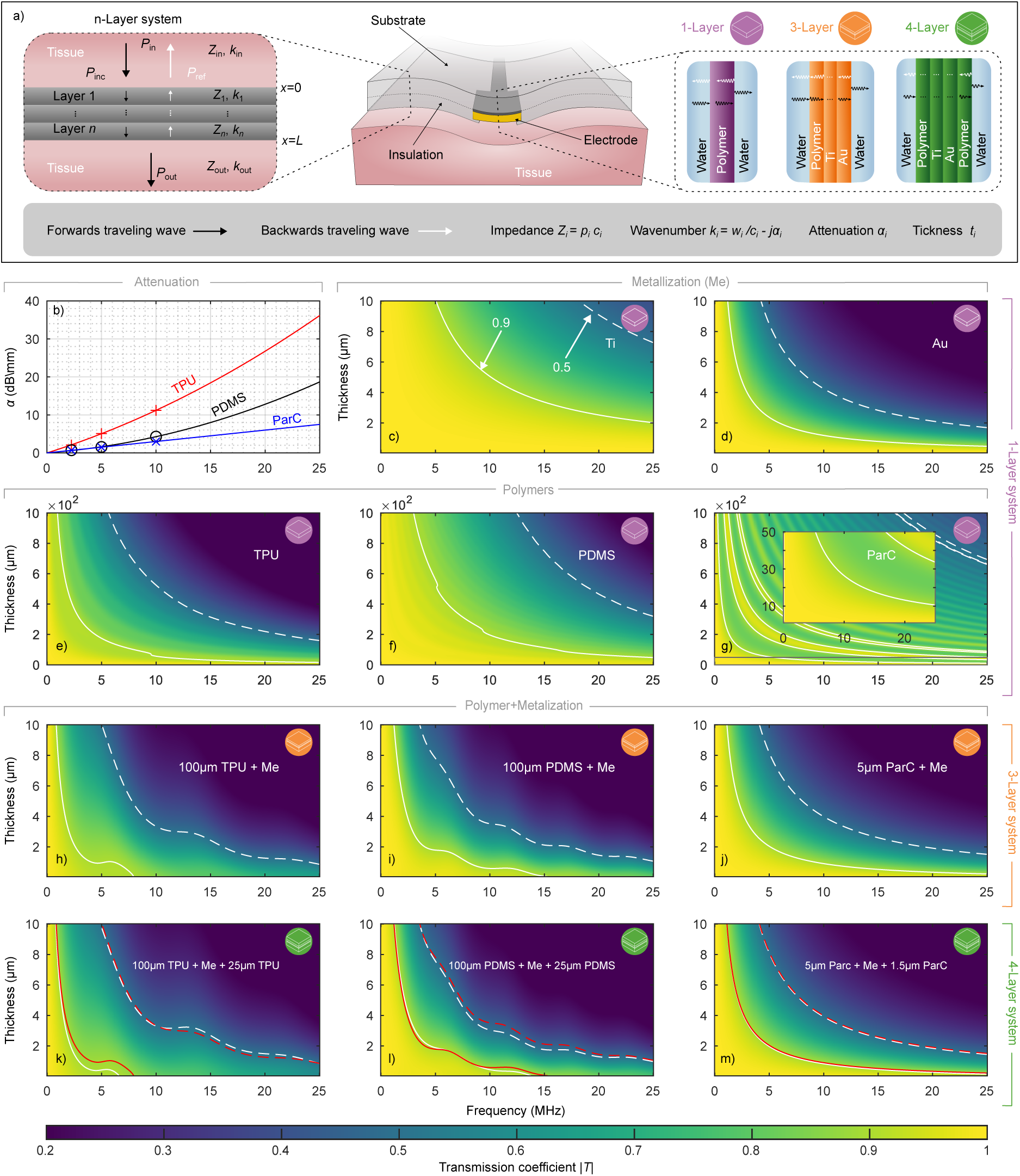
Structure and acoustic modeling of polymer-based electrodes. a) Schematic representation of the multilayer acoustic model used to simulate ultrasound propagation through neural interfaces. b) Frequency-dependent acoustic attenuation coefficients for selected polymers (TPU, PDMS, ParC), compiled from literature^99,100^. Solid lines denote quadratic fits to the reported data. c–g), Numerical simulations of the ultrasound transmission coefficient as a function of frequency and thickness for single-layer systems of titanium (c), gold (d), TPU (e), PDMS (f), and ParC (g). Insets in (g) for adapted axis. h–j), Simulated transmission maps for three-layer stacks comprising a polymer substrate (100 µm TPU, PDMS, or 5 µm ParC), with a 50 nm Ti adhesion layer and variable-thickness Au metallization (Me: 50 nm Ti + variable-thickness Au). k–m), Transmission maps for four-layer systems incorporating an additional insulation layer (25 µm TPU, PDMS, or 1.5 µm ParC). Solid contours indicate 0.9 transmission, and dashed contours indicate 0.5 transmission. Dashed red contours indicate the transmission threshold of the corresponding three-layer configurations, for direct comparison.

To assess the individual contributions of each material to acoustic transmission, we first simulated single-layer structures composed of common polymers used for substrates or insulation, and metals typically employed for tracks or electrodes (Figs. 3c–3g, reference thresholds of 0.9 and 0.5 are indicated in the graphs). Additional material simulations can be found in Supplementary Fig. 1. Model parameters are summarized in Fig. 2b. Metallization types and thicknesses were initially simulated to identify configurations with minimal acoustic loss. Figs. 3c and 3d show the transmission coefficient as a function of frequency for single-layer of Ti and Au, respectively. Both metals exhibit non-linear transmission losses that increase with frequency and thickness. As intrinsic attenuation in metals was not included in the model, the observed transmission losses arise solely from acoustic impedance mismatches at the interfaces, which result in partial wave reflections.

The results reveal stark differences between the two materials. Ti displays relatively favourable transmission characteristics across the simulated frequency range. Notably, a 2 µm Ti layer maintains over 90 % acoustic transmission, highlighting its suitability for high-frequency ultrasound applications. In contrast, Au shows substantially poorer transmission under identical conditions, dropping below 50 %, due to its greater acoustic impedance and resulting acoustic mismatch. To preserve favourable acoustic transparency at higher frequencies, Au layers must be thinner, typically less than 1 µm.

For the polymer, substrates we assessed TPU, PDMS and ParC. Figs. 3e–3g present the transmission coefficients for the polymer layers, incorporating material-specific attenuation Fig. 3b. As with metals, transmission through polymers decreases non-linearly with increasing frequency and thickness. PDMS shows slightly higher transmission than TPU, primarily due to its lower attenuation. In both materials, attenuation becomes the dominant loss mechanism at thicknesses exceeding a few hundred micrometres, particularly at higher frequencies. ParC, which has a comparatively low attenuation coefficient, reveals distinct transmission peaks associated with acoustic resonances, occurring at conditions satisfying *t_i_* = *nλ/*2, where *n* = 0, 1, 2, These resonances are more pronounced in ParC than in PDMS or TPU, due to its different attenuation profile. However, ParC achieves similar transmission only at substantially smaller thicknesses, nearly one order of magnitude thinner, due to its higher acoustic impedance mismatch with the surrounding media. Finally, as seen in (Figs. 3c–3g), *T* approaches one as layer thickness *t_n_*approaches zero. This asymptotic behaviour provides a baseline for interpreting the influence of material properties and layer geometry on ultrasound propagation.

Building on the single-layer analysis, we next evaluated more complex structures representative of practical device architectures. Specifically, we modelled an open electrode stack consisting of a polymer substrate, a 50 nm Ti adhesion layer, and a variable-thickness Au layer (up to 10 µm). Transmission performance for stacks based on 100 µm TPU, 100 µm PDMS, and 5 µm ParC is shown in Figs. 3h–3j. In all cases, the Au thickness primarily limits transmission due to reflection losses, while the polymer attenuation adds secondary losses. This effect is further illustrated in Figs. 3k–3m, where we added an additional polymer layer-25 µm TPU, 25 µm PDMS, or 1.5 µm ParC-representing the insulation layer of a neural interface. While the addition of an insulation layer introduces an extra acoustic interface and some attenuation, its impact remains minimal, with effects primarily emerging at higher frequencies. In certain configurations, it can even enhance transmission, as shown in Fig. 3k, where the 0.5 contour exhibits slight improvements. Increasing the insulation thickness to match the substrate (see Supplementary Fig. 2) leads to only modest reductions at higher frequencies. A similar trend is observed when the total polymer thickness is conserved, and the metallization is embedded symmetrically between two polymer layers (see Supplementary Fig. 2). For parylene, these adjustments have a negligible impact. Resonance features remain evident along the 0.5 and 0.9 contours for TPU and PDMS, indicative of constructive interference within the multilayer stack. Overall, the insulation layer exerts only a minor influence on transmission, with optimal performance achieved when its thickness is kept to a minimum.

These results highlight that thin, impedance-matched multilayer structures can enhance ultrasound transmission, particularly at longer wavelengths, and that resonance conditions can be exploited to improve transmission at shorter wavelengths.

### Experimental validation of the numerical simulations

To validate the accuracy of our numerical framework, we conducted ultrasound transmission experiments using a water tank setup with multilayer configurations at selected thicknesses matching simulated scenarios. These included single-layer polymer sheets to represent substrate and insulation materials, a three-layer stack (polymer + 50 nm Ti + variable-thickness Au) for exposed electrode structures, and a four-layer configuration for embedded or encapsulated tracks. A single-element, unfocused ultrasound transducer generated the acoustic field, which was recorded by a needle hydrophone placed opposite the transducer (Fig. 4a). Transmission was quantified by comparing ultrasound pressure amplitudes with and without samples positioned between the transducer and hydrophone, across a frequency range of 1.25-25 MHz. We further assessed the compatibility of the proposed framework with FUS by analysing the influence of a ParC-embedded metal multilayer structure (5 µm ParC + 50 nm Ti + 300 nm Au + 1.5 µm ParC), specifically designed to replicate realistic material properties of a neural interface, hereafter referred to as the Neural Electrode Phantom (NEP), and a patterned electrode array (2 µm ParC + 5 nm Ti + 80 nm Au + 1.5 µm ParC) on the transmission coefficient and beam profile of a 16.35 MHz FUS transducer. Additional methodological details are provided in the Methods section.

**Figure 4.**
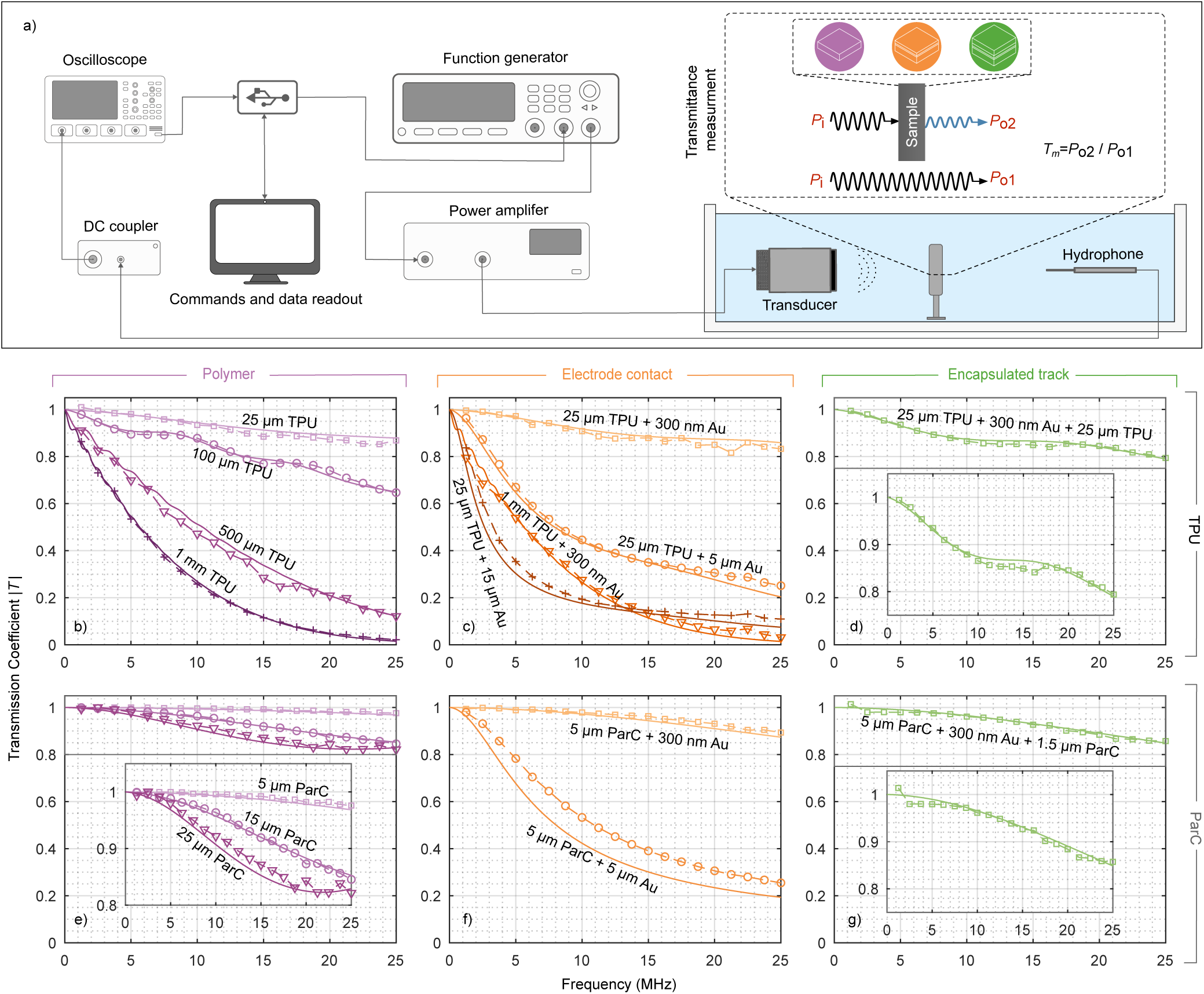
Experimental setup for ultrasound transmission and comparison with numerical model predictions. a) Schematic of the experimental setup used to quantify ultrasonic pressure transmission across multilayer samples. A function generator drives a single-element transducer via a power amplifier, producing an acoustic field in a water tank. The transmitted pressure is captured by a needle hydrophone placed opposite to the transducer. Transmittance (*T* _m_) is calculated by comparing the measured pressure with (*P*_o1_) and without (*P*_o2_) the sample in the acoustic path. Water serves as a tissue-mimicking medium due to its comparable acoustic properties. b-g) Comparison between numerical simulations (solid lines) and experimental measurements (dashed lines with markers) of the transmission coefficient (*T* ) for TPU and ParC-based systems. One-layer configurations using TPU (b) and ParC (e) at various polymer thicknesses. Three-layer configurations, representing an electrode contact consisting of 50 nm Ti and varying Au and polymer thicknesses, for TPU (c) and ParC (f). Four-layer configuration, representing an encapsulated track using a TPU (d) stack comprising a 25 µm base layer, 50 nm Ti, 300 nm Au, and a 25 µm insulation layer. Four-layer configuration, representing an encapsulated track using ParC (g) with a 5 µm substrate, 50 nm Ti, 300 nm Au, and a 1.5 µm insulation layer. Insets show zoomed-in views with adapted axes to enhance clarity.

Across all materials and thicknesses, we observed excellent agreement between simulations (*T* _s_) and measurements (*T* _m_), with transmission decreasing non-linearly at higher frequencies and layer thickness. In the 25 µm TPU sample (Fig. 4b), high transmission was maintained across the entire frequency range, remaining close to 90 % (T_m_∼0.88 at 25 MHz). For the 100 µm sample, resonance features were clearly observed, with the first peak occurring near ∼8.75 MHz, consistent with the expected condition *f* = *c/*2*t*. However, these resonances are substantially damped due to intrinsic material attenuation. They are consequently absent in the thicker TPU samples, as well as in the 25 µm sample, where the first resonance would be expected beyond the evaluated frequency range (∼35 MHz). Experimental results for ParC (Fig. 4e) further confirm the simulations, showing preserved transmission of more than 80 % across the tested thicknesses and frequencies. In the three-layer system Figs. 4c and 4f, the 25 µm TPU sample with a 300 nm Au layer maintained over 80 % transmission even at 25 MHz. A direct comparison between 25 µm and 5 µm ParC configurations highlights a key insight: similar transmission performance can be achieved with markedly different materials and geometries, underscoring the importance of acoustic impedance matching and polymer selection for device optimization.

Regardless of the polymer type, the transmission coefficient for thin polymer layers is predominantly dictated by the metal layer. Conversely, high transmission can be retained when the metal layer is thin, provided the polymer thickness remains moderate. As the polymer thickness increases, attenuation within the polymer becomes more pronounced, leading to a substantial reduction in transmission. This trend is clearly demonstrated in Fig. 4c, where the 1 mm thick TPU layer with a 300 nm Au shows a noticeable drop in transmittance. The numerical and experimental results for the four-layer system Figs. 4d and 4g similarly show that metallization dominates the transmitting behaviour, with only minor additional loss from the added insulating layer. Fig. 4d also shows the local resonance at approximately 15 MHz. Finally, the FUS measurements showed subtle differences in the beam profile and peak pressure between the conditions with and without the sample, although these were not statistically significant. Results of the analysis are presented in Supplementary Fig. 3 and Supplementary Fig. 4. As predicted by prior simulations (Fig. 4g), an acoustic loss of approximately 8 % was expected for the NEP. However, experimental peak amplitude measurements show a loss of around 4 %, suggesting that wave propagation at shallower incidence angles reduces reflection-related losses. The measured difference in experimental peak amplitude for the electrode array is even less than 1 % (Supplementary Fig. 4f).

### Compatibility with functional ultrasound imaging in a flow phantom

To determine whether functional ultrasound (fUS) signals can be effectively recorded through a neural interface, we first developed an ultrasound flow phantom. Briefly, a silicon tube with an inner diameter of 200 µm was embedded in gelatin using the proposed protocol, detailed in the Methods section (Fig. 5a), and a blood-mimicking fluid (BMF) was pumped through. The recording principle of fUSI is shown in Fig. 5b and detailed in the Methods section. Tilted ultrasound plane waves are generated using an 18 MHz transducer array. The recorded backscattered signals are then utilised for the reconstruction of compound images. Following coherent summation and clutter filtering, ensembles of compound images generate the power Doppler images. The mean image, constructed of all power Doppler images, is referred to as the mean power Doppler image (MPDI). We compared recordings obtained without the NEP (w/o) to those with the NEP (w/) placed on top of the phantom (Fig. 5c). The power Doppler signal intensity decreased with increasing depth and shows a statistically significant decrease when the NEP was present (Fig. 5d). Moreover, the signal-to-noise ratio (SNR) decreased with depth of the imaging plane. However, the presence of the NEP did not result in a significant change of the SNR compared to the control (Fig. 5e) at the imaging depth of 6 mm. The results in Fig. 5c indicate the mean noise level and standard deviation within the region-of-interest (ROI) of the MPDI, while Fig. 5e represents the mean and standard deviation of individual acquisitions (180 frames). Examination of the full width at half maximum (FWHM) of the in-plane point-spread function (PSF) reveals a more distinct influence of the NEP (Fig. 5f). The lateral and axial FWHM deteriorate with imaging depth, which is to be expected due to the attenuation of the acoustic fields. This leads to the smearing of the deeper-lying target due to a decrease in the power Doppler intensity in the ROI and an increase in the noise level. However, in the presence of the NEP, a decrease in the FWHM can be observed in the lateral and axial directions at each imaging depth, compared to the respective FWHM without the NEP. This can be explained by the fact that the neural interface has minimal impact on the noise level, but the signals in the ROI are reduced. As a result, the PSF appears slightly narrower, which is not necessarily associated with a higher resolution, but rather indicates a loss of signal (see Supplementary Fig. 5). Although significant statistical differences between the two conditions are revealed, the qualitative evaluation of the power Doppler images suggests that these differences may not be practically meaningful. Despite the statistical significance, the Doppler signals observed in the images remain consistent and reliable across both conditions. Upon visual inspection, the strength and clarity of the Doppler signals do not exhibit any substantial or perceptible variation that would indicate a meaningful difference in the power Doppler signal. This suggests that, while the statistical test identifies a difference, the functional impact of this difference may be negligible from a practical perspective. In essence, while the data may suggest some variation in the measurements between conditions, the visual analysis of the Doppler signals indicates that these differences are unlikely to affect the overall interpretation of the imaging results. In addition to the NEP measurements, we evaluated fUS recordings acquired both without an electrode array and with the array placed on the ultrasound phantom. As in the NEP experiments, small measurable differences were observed. However, visual inspection of the reconstructed Doppler images showed no substantial or practically meaningful changes. The intensity, SNR, and overall appearance of the Doppler signals remained consistent across both conditions, indicating that the electrode array, given the metallization thicknesses and pattern dimensions used here, does not noticeably impact the imaging outcome (Supplementary Fig. 6).

**Figure 5.**
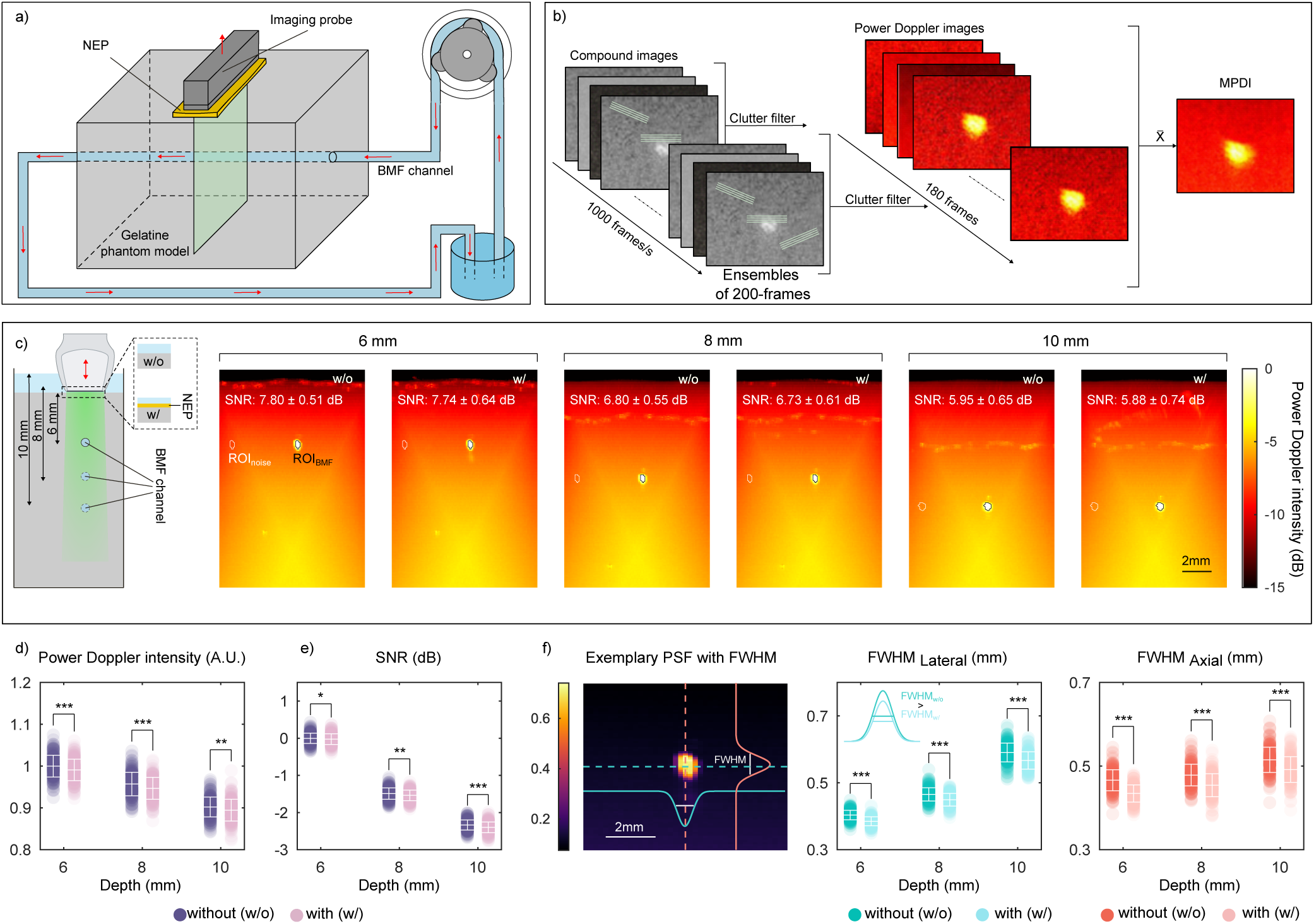
Compatibility of functional ultrasound imaging with neural interface using a flow phantom model. a) Schematic of the flow phantom. b) Schematic of the principles of fUS imaging. c) Schematic of the data recording principle with the US probe placed at three imaging depths (6, 8, and 10 mm) above the BMF and MPDI of the blood flow phantom, acquired without (w/o) and with (w/) the NEP. The SNR within the ROI is reported as mean ± standard deviation (SD). d-e) Power Doppler intensity (d) and SNR (e) of ROI with and without the NEP, at various imaging depths. f) Characterization of lateral and axial resolution. Exemplary PSF with FWHM in lateral and axial directions across the flow channel cross-section. Data are presented as mean ±SD; dots represent individual acquisitions, n = 180 acquisitions per group. (All values were normalised against the control (MPDI, no NEP at 6 mm imaging depth). *P > 0.05, **P < 0.01, and ***P < 0.001)

Lastly, we recorded electrical signals from an electrode array during power Doppler imaging under four configurations (see Supplementary Fig. 8). Recorded artifacts matching the duration of the ultrasound pulses were detected in most scenarios, indicating interference from the US probe. When the NEP was used as additional shielding, no artifacts were observed, confirming electromagnetic coupling as the source and demonstrating that proper shielding prevents this interference. Although the shielding may introduce slight additional losses, these are negligible in the present experiments, given their minimal effect on overall transmission (see Supplementary Fig. 7).

### Functional ultrasound imaging through ultrasound-transparent neural interfaces in vivo

We further investigated the feasibility of detecting functional brain signals with ultrasound through the NEP in awake mice. The experimental design is presented in Fig. 6a. Briefly, to trigger a neuronal response in the mouse brain, we exposed the eyes of the head-fixed mice to a visual stimulation protocol. This protocol was designed to evoke responses in brain regions that are part of the visual system, namely the visual cortex, the superior colliculus (SC) and the lateral geniculate nucleus. The ultrasonic probe was positioned over a coronal plane, presenting these vision-involved structures. We conducted fUSI in two mice across two sessions: one prior to, and one following, placement of the NEP on the cranial window surface. In each session, a baseline recording without visual stimulation (rest) was performed first, followed by recordings during stimulus presentation (task). More details can be found in the Methods section. Data from one mouse were excluded due to excessive motion artifacts during recording. Fig. 6b(i) and Fig. 6c(i) illustrate the mean functional ultrasound image (MPDI), generated by averaging 5752 frames acquired from the mouse brain in the absence and presence of the NEP. In vivo results showed that the mean power Doppler intensity (Fig. 6d) within the subcortex region (SCR) and deep brain region (DBR) decreases when the NEP is present by 25 % and 29 %, respectively. The SNR (Fig. 6e) in the SCR decreased by 1.3 dB and for the DBR by 1.6 dB. In both experimental conditions, the general linear model (GLM), used to generate the activation map of Fig. 6, identified functionally active regions within the SC, characterised by positive signal modulation during stimulation across individual voxels (Fig. 6b(ii) and Fig. 6c(ii)). However, in the control recording, fewer voxels showed significant activation (*η*^2^ > 0.1) within the SC region. As expected, presentation of the visual stimulus evoked a pronounced response in the SC region (ROI I), clearly observable during the task block in Fig. 6b(iii) and Fig. 6c(iii). Although no statistically significant changes were detected in ROI I when comparing the two conditions over the full experiment duration (P > 0.05, two-sided Student’s test), a decrease of 6.36 % in average activity during the task block was observed, suggesting a modest, though non-significant (P = 0.16, two-sided Student’s test), signal decrease. This observation is further supported when considering the temporal signal-to-noise ratio tSNR (Fig. 6f), which shows a reduction of 0.06 dB in the active region when comparing the two conditions. The tSNR in the non-active region (ROI II) decreases by 0.02 dB. The signal within ROI II (Fig. 6b(iii) and Fig. 6c(iii)) also remained stable across the entire recording with no significant differences observed between conditions (P > 0.05, two-sided Student’s test). All values were normalised against the control scenario (no NEP, SCR or ROI I). To assess the consistency of structural features across the two experimental conditions, we quantified the similarity between paired images using the Structural Similarity Index Measure (SSIM). Overall, the SSIM values indicated a high degree of correspondence between conditions, confirming the reliability of the imaging approach under varying acquisition parameters. The structural similarity index measure (SSIM) map, comparing the two MPDI (Fig. 6g), displays values predominantly above 0.75, with the lowest similarity localised in regions of elevated blood flow. Correspondingly, the SSIM histogram (Fig. 6h) reveals a strong structural resemblance between the two conditions, with indices consistently exceeding 0.7 and the majority surpassing 0.9. These findings indicate that the presence of the NEP introduces minimal interference and affirm the reproducibility of functional ultrasound imaging under these conditions.

**Figure 6.**
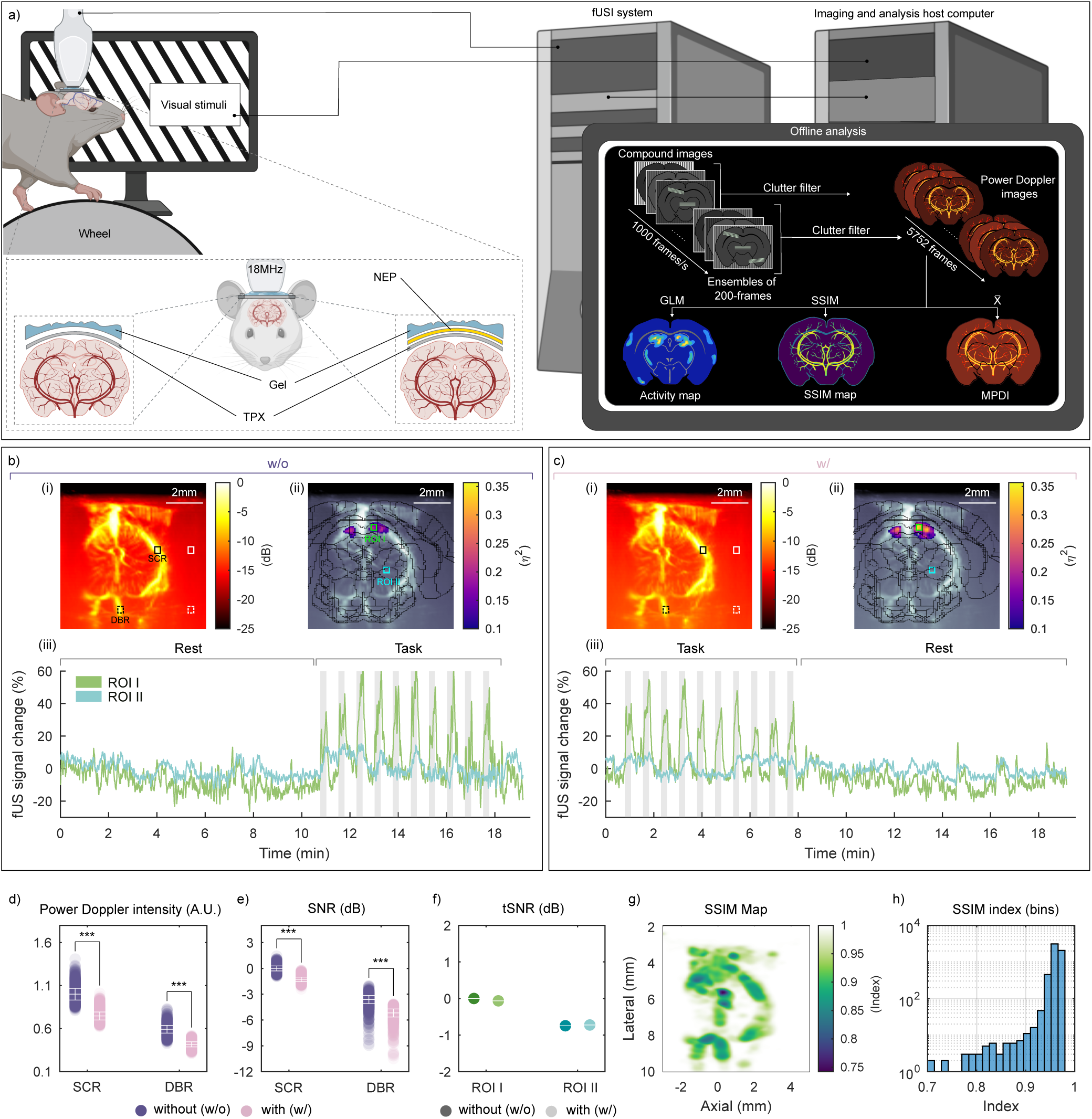
Functional ultrasound Doppler imaging of mice brain through a flexible neural interface. a) Schematic of the experimental setup for fUS brain imaging through a coronal plane during visual stimulations^101^. (b-c) mean power Doppler image (MPDI) of mouse brain (i), activation map (ii) and mean fUS signal (iii) without (w/o) interface b) and with (w/) c) interface placed above the cranial window. Squares indicating ROIs. SCR and DBR in (i) for SNR and power Doppler intensity in (d) and (e), respectively. ROI I and ROI II in (ii) for the temporal signal in (iii). d) power Doppler intensity of the SCR and DBR for each condition. Data are presented as mean ±SD with arbitrary units (A.U.). e) SNR of ROIs. Data are presented as mean ±SD. f) tSNR of Doppler signals of ROIs (ROI I = SC region with high effect size (green) and ROI II = deep brain region with low effect size (blue)). g) SSIM map of the two MPDI with and without NEP. h) Histogram of the overall SSIM index comparing single frames of fUSI recordings with NEP against the control. All values were normalised against control (MPDI, no NEP, SCR or ROI I). Dots represent individual acquisitions, n = 5752 acquisitions per group. ***P < 0.001)

## Discussion

In this study, we introduce a comprehensive framework for quantifying the ultrasound transmission properties of neural interfaces, aiming to support their integration into multimodal platforms that combine electrophysiological recording with ultrasound imaging. Prior to in vivo testing, we systematically evaluated the acoustic performance of relevant materials using both numerical simulations and in vitro experiments. Building on foundational studies of ultrasound propagation through multilayered systems^33^, our work advances the understanding of how soft, flexible neural interfaces can be optimised for acoustic transparency. We developed a theoretical model that captures the key transmission characteristics of multilayer polymer-metal composites commonly used in bioelectronic devices. Notably, the numerical model assumes idealised conditions, specifically perpendicular continuous plane wave propagation at a single frequency. In contrast, ultrasound under realistic conditions is inherently three-dimensional and temporally pulsed, encompassing a broad spectrum of frequency components. While the assumption of ultrasound waves travelling perpendicular to the array surface provides a reasonable approximation for micro-electrocorticographic (µECoG) array architectures, the proposed model is also applicable to scenarios involving angled wave propagation^34^, which are more likely in the case of penetrating probes. This is because the present study focused on capturing the most conservative case, where incident waves are perpendicular to the interface, resulting in maximal acoustic reflection due to impedance mismatches at material boundaries^35^. In contrast, wave propagation parallel to the neural interface represents a more specific case that may involve complex transmission dynamics and requires further investigation.

Additionally, while the model accounts for attenuation within the polymer layers, it does not explicitly include energy losses due to metallization, effects such as surface scattering or dissipation in the metal layers^36^. These omissions likely contribute to minor deviations between simulations and measurement at higher frequencies. Nonetheless, the overall agreement between model predictions and empirical data supports the framework’s utility as a predictive tool for designing acoustically transparent neural interfaces. Our findings also emphasise that materials with acoustic properties more closely matched to the surrounding medium tend to offer higher transmission efficiency. Our results show high ultrasound transmission through metallization layers several hundred nanometers thick. Similar or even thinner layers have been used in practice for neural electrodes^37–44^. However, trade-offs inherent to interface design must still be carefully considered. We were able to show through simulation that a 2 µm thick Ti layer can preserve 90 % transmission at 25 MHz, but may not be the preferred electrode material due to its higher electrical impedance^45^. Similarly, thicker layers can exhibit favourable resonance-enhanced transmission, but at the cost of increased stiffness^46^ and frequency specificity condition, as shown for ParC (Fig. 3 and Fig. 4). Our findings also confirm that thin insulation layers contribute minimally to overall losses, and that nanometer-scale metallization is sufficient to maintain both practical electrical impedance^47–50^ and high acoustic transmission. We validated our approach through fUSI in phantoms and awake mice using an appropriately designed multilayer structure as a neural interface-mimicking structure. Consistent with simulation results, the presence of the interface did not appreciably distort the Doppler signal. Despite the presence of the interface, fUSI revealed robust activation of the SC. While a measurable influence was detected using a deliberately structured NEP, no imaging artifacts were apparent in the resulting Doppler data. This supports the conclusion that the acoustic footprint of the interface can be minimal under imaging conditions. In contrast to the findings in^32^, which indicated a smaller effect size for highly attenuating or reflective materials within the cranial window, we observed a larger effect size in our setup. These findings suggest that the variability introduced by the experimental conditions exceeds the influence of the electrode itself. We note, however, that the absence of a consistent effect in the GLM output should not be taken to imply the absence of an interface-related influence. Rather, the effect size may be small relative to the overall variability across sessions, and potentially masked by the limited sample size (N=2). Future studies involving larger animal sample sizes will be essential for disambiguating subtle interface-related effects and for improving estimates of their impact on fUSI outcomes, particularly when targeting smaller or less robustly activated structures. In parallel, advances in imaging reconstruction methods, particularly aberration correction strategies that account for sound speed heterogeneities across layered tissue structures^51^, have markedly improved the quality of non-invasive transcranial fUS imaging. While originally developed to address skull-induced distortions, this modelling framework could, in principle, be extended to correct for aberrations introduced by implanted neural interfaces. Together, these complementary developments point toward the realisation of integrated, multimodal neurotechnologies, in which both electrode design and beamforming strategies are co-optimised to maintain high imaging fidelity and signal integrity. Given the broad use and recent advantages of ultrasound techniques in the medical field^18,51–53^, the potential translational prospects for acoustically transparent neural electrodes are exciting. The combination of ultrasound technologies and electrophysiological techniques could not only enable concurrent recordings in both domains and therefore would allow capturing fast transitions in brain states with electrophysiology while observing slow vascular changes using fUSI, linking local neural activity to hemodynamics^31^ and brain-wide mapping of local activity. Moreover, emerging materials such as graphene, PEDOT:PSS, and hydrogels are being explored as alternatives to traditional materials for neural electrodes, each offering distinct acoustic advantages. Graphene is promising due to its atomically thin structure, which helps reduce acoustic losses. PEDOT:PSS combines moderate acoustic properties with the benefit of being deposited in thin layers, improving acoustic transparency. Hydrogels, with their tissue-matched acoustic impedance, offer excellent coupling with biological tissue. These properties make each material individually attractive for enhancing the acoustic performance of neural electrodes. A qualitative survey of these candidate materials, along with key acoustic and mechanical parameters, is provided in Supplementary Note 2 and Supplementary Tab. 1. Irrespective of the substrate material or its degree of ultrasonic transparency, the interaction between ultrasound waves, biological tissue, and implanted interfaces presents technical challenges, particularly at higher acoustic intensities. These challenges are especially relevant for ultrasound neuromodulation, where focused ultrasound enables localised modulation of neural activity while electrode arrays concurrently record the resulting responses. A principal concern in such contexts is the acoustoelectric effect, in which pressure fluctuations from the ultrasound field induce local modulations of the electric field at the electrode–tissue interface, potentially generating artifacts in the recorded signals.^54–58^. This effect is especially pronounced for high pressure and in the presence of externally applied electric fields, as during concurrent stimulation and recording. In contrast, fUSI operates at considerably lower acoustic intensities, reducing the likelihood of appreciable acoustoelectric interference. Moreover, the resulting artifacts are typically confined to ultrasonic frequencies and therefore well outside the frequency range of the neuronal signals^59^, making them likely to affect low-frequency electrophysiological data. Nevertheless, as no electrophysiological experiments were performed in this study, the impact of acoustoelectric coupling during concurrent fUSI and electrophysiology remains to be evaluated. Additional sources of interference include mechanical displacement of the neural electrode, such as that caused by acoustic radiation forces or bulk vibrations, as well as electromagnetic coupling between the recording system and the high-voltage drivers used to operate ultrasound transducers. While electromagnetic coupling is a common cause of signal corruption, ranging from transient disturbances to amplifier saturation, due to the physical proximity between sensitive recording electronics and power-intensive ultrasound hardware, mechanical effects are most pronounced under high-intensity ultrasound exposure^60–62^. To mitigate electrical interference, careful system design is required. Specifically, robust shielding, proper grounding, and spatial decoupling of stimulation and recording components. These measures are critical for the reliable integration of ultrasound technologies with electrophysiological systems. Further details and representative data illustrating electrical interference, along with mitigation strategies, are provided in the Supplementary Fig. 8. While our study specifically focuses on neural electrodes compatible with fUSI, where the simplifying assumptions employed in the model are sufficient to capture realistic transmission behaviour, we further believe that this modelling approach can be extended to applications involving FUS. To support this, we performed additional measurements using a FUS transducer and conducted beam analysis, presented in the Supplementary Fig. 3. Simultaneously, we performed the same beam analysis with the complete electrode array in place, which revealed no significant changes in the ultrasound field (Supplementary Fig. 4.) These findings support the assumption that reflection decreases at lower incidence angles and validate the hypothesis that the proposed model provides an upper boundary framework for selecting materials and compositions suitable for acoustically transparent neural electrodes. It is important to note, however, that the model is not intended to capture detailed field characteristics. Rather, it serves as a design-oriented framework for identifying suitable materials and compositions for acoustically transparent neural electrodes. The presented model is limited in handling 2D and 3D problems because it assumes planar, laterally infinite layers and cannot account for curved interfaces, localised inhomogeneities, or arbitrary geometries. It also becomes numerically unstable in thick or highly layered structures and cannot accurately model wave scattering or complex boundary conditions^63,64^. For cases requiring accurate resolution of realistic acoustic field distributions, full numerical methods such as FEM or k-Wave are recommended. However, these methods can be computationally demanding, as resolving sub-micrometre features often requires fine spatial discretisation and significant computational resources. Readers interested in such simulations are referred to^65–68^. Nevertheless, our findings suggest that focused ultrasound stimulation could be paired with ultrasound transparent electrode arrays to enable simultaneous modulation and recording. Although this integration remains to be experimentally validated, the conceptual framework we provide lays the groundwork for such integration. In summary, we demonstrate that flexible neural interfaces can achieve high ultrasound transmission across clinically relevant frequencies through careful engineering of material properties and structural design. Key design strategies include thin to moderately thick polymer layers (ranging from a few to tens of micrometres), metal layers that are deeply subwavelength thin, and impedance-matched stacks that minimise reflection and signal attenuation. Metal layers that are small relative to the wavelength (10^−3^-10^−2^ *λ* ) remain largely transparent, exerting minimal influence on the ultrasound field. Although the impact of more complex electrode features, such as line–space architectures with metallization approaching a non-negligible fraction of the wavelength, warrants systematic investigation in future studies, our results indicate that the electrode designs proposed here are effectively ultrasound transparent. Given the growing clinical and research use of ultrasound-based techniques, acoustically transparent neural interfaces represent a promising route toward advanced multimodal neurotechnologies that unify electrical and hemodynamic readouts and, potentially, therapy delivery in a single platform.

## Methods

### Acoustic transmission through layered systems

Under the assumption of a one-dimensional system with *n* layers with thickness *t_i_*, between two semi-infinite media, the acoustic transmission coefficient for a longitudinal plane wave travelling across a distance *x* (Fig. 3a) at a frequency *f* , can be obtained by applying the TMM^34^. Assuming continuity of pressure and particle velocity at each boundary, the total transfer matrix M relates the sound pressure and acoustic particle velocity at the left boundary (input side, *x*=0) of the structure to that at the right boundary (output side, *x*=L) of the structure:

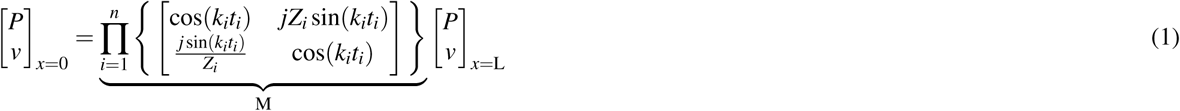

where *Z* = *ρc* is the characteristic acoustic impedance, 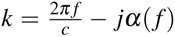 is the complex wavenumber with *α*( *f* ) as the frequency-dependent acoustic attenuation coefficient, *ρ* the density and *c* the sound speed of the material. The pressure and velocity on the input side (*x*=0) can be written as

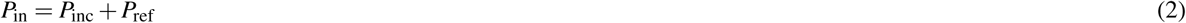

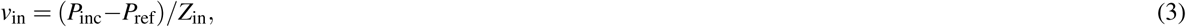

and velocity on the output side (*x*=L) can be written as

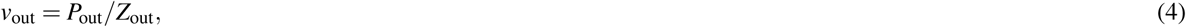

where, *P*_inc_ and *P*_ref_ are the pressure amplitude of the incident and reflected wave on the input side and *P*_out_ the pressure amplitude on the output side. Thus, Eq. (1) can be written as

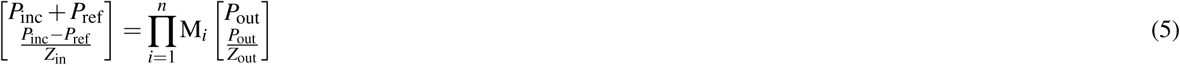

The results are then obtained by the expansion of the matrix for *n* layers between two semi-infinite layers and the elimination of *P*_ref_. The transmission coefficient can be found by 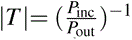, using a custom MATLAB script (see Supplementary Note 1 for analytical solutions). We analysed the acoustic properties of the selected materials by simulating the acoustic transmission coefficient |*T* | through polymer and metal layers of varying thickness. For the simulations, we selected representative thicknesses to span a broad range from nm to mm scale to cover a large area of material thicknesses. Additional stratified material stacks data can be found in Supplementary Fig. 2. The material properties used are listed in Fig. 2b and are based on previous literature.

### Transmittance measurement setup and procedure

The experiments were conducted in a 580 x 450 x 300 mm^3^ water tank. Deionised water was used as the intermediate medium as its density, sound velocity, and sound impedance are near those of soft tissue. An acoustic-damping material was attached to the sides of the tank to minimise acoustic reflections. An ultrasonic transducer (V309-SU, Olympus, Japan) with a diameter of 0.5 inch (12.7 mm) was driven by an arbitrary function generator (33622A, Keysight, USA) and a 50 W RF power amplifier (350L, Electronics & Innovation, USA). An ultrasonic pulse train comprising modulated sine waves (15 µs duration, 4 µs rise and fall times) was employed, sweeping frequencies from 1.25 MHz to 25 MHz in 1.25 MHz increments. The radiated ultrasonic signal passed through the fabricated samples, aligned and positioned 10 mm away from the transducer surface. To perform measurements of through-transmitted energy, we employed a needle hydrophone (NH1000, Precision Acoustics, UK) with a diameter of 1 mm at a distance of 35 mm away from the transducer. The hydrophone was coupled with a submersible preamplifier and DC coupler. The samples were tested one at a time. By measuring the pressure with and without the sample placed between the transducer and the hydrophone, transmittance can be calculated by comparing the pressure collected through the sample (*P*_o2_) and without the sample in the path (*P*_o1_). A custom-made 3D print was used to align the transducer, sample and hydrophone. The data acquired from the hydrophone were recorded using a digital storage oscilloscope (RTA4004, Rohde & Schwarz, Germany). All applied and monitored signals were time-synced between the function generator and oscilloscope with trigger inputs. The monitored signals were averaged over 100 cycles and recorded with 1.25 GSa*/*s. The logged signals were streamed from the data acquisition hardware to a workstation PC using custom Python code to control the triggers, channels, and function generator output. The compiled C code, which called the oscilloscope and function generator commands, was in turn called within a Python wrapper. The data were then subsequently processed using MATLAB.

### Focused ultrasound transmittance measurement setup and procedure

The experiment was performed following a protocol analogous to that used for transmittance measurements. All measurements were carried out in a water tank measuring 580 x 450 x 300 mm^3^, filled with deionised, degassed water. An acoustic-damping material was attached to the sides of the tank to minimise acoustic reflections. A custom-ordered focused ultrasonic transducer (PA2729, Precision Acoustics, UK) with a centre frequency of 16.35 MHz and a diameter of 19 mm was driven by an arbitrary function generator (33622A, Keysight, USA) and a 50 W RF power amplifier (350L, Electronics & Innovation, USA). An ultrasonic pulse train comprising modulated sine waves (20 µs duration, 4 µs rise and fall times) was employed. To perform measurements of through-transmitted energy we positioned a needle hydrophone (NH1000, Precision Acoustics, UK) with a diameter of 75 µm in the focal spot (at a distance of 20.41 mm from the transducer surface). The hydrophone was mounted on a motorized XYZ stage, enabling 3D mapping of the ultrasound beam by scanning across the field. The hydrophone was coupled with a submersible preamplifier and DC coupler. Lateral 2D scans were performed alternately with and without the probe placed between the transducer and hydrophone. To account for small setup instabilities or movements, each scan session was repeated four times. The sample was fixed in place using a custom-made 3D-printed frame, which was mounted on a custom-built sample holder. The NEP was used as sample (5 µm ParC substrate, 300 nm Au with a 50 nm Ti adhesion layer, and 1.5 µm ParC insulation). Gaussian functions were fitted to the beam profiles along both the axial and lateral axes through the centre of the beam. Peak pressure and FWHM values were extracted. All measurements were normalised to the mean intensity of the control condition (i.e., no NEP present). Additionally, analogous measurements were performed using the electrode array with the insulated tracks placed within the ultrasound field. The electrode array fabrication is detailed in reference^69^. Briefly, 2 µm thick, ultra-flexible ParC substrate is deposited by chemical vapor deposition (CVD), subsequently, 80 nm thick gold interconnects with a 5 nm titanium adhesion layer are photolithographically patterned, followed by deposition of a 1.5 µm thick insulating ParC layer over the metal lines using a silane adhesion promoter.

### Sample fabrication for transmittance measurements

For the experiments free-standing material films, different polymer (TPU, ParC) thicknesses and metals (Au, Ti) were chosen to evaluate the numerical simulations over a wide range of parameters while maintaining mechanical stability and ease of handling. For both TPU (Platilon 4201 AU, Covestro AG, Germany) and ParC (type C, Galentis S.r.l., Italy), three sample sets were fabricated using standard microfabrication steps. For ParC, the first step comprised in the deposition of 5 µm, 15 µm or 25 µm-thick layers on 22 mm x 44 mm microscope glass slides through a CVD process (Comelec SA, Switzerland). Prior to this, the glass slides were cleaned in deionised water and isopropanol. Subsequently, for the three-layer samples, 300 nm, 5 µm or 15 µm-thick Au metallization with a 50 nm Ti adhesion layer were deposited in a sputter system (Creavac, Germany) on 5 µm ParC. For the four-layer system, one sample was prepared by depositing 5 µm ParC, 300 nm Au with a 50 nm Ti adhesion layer, and another 1.5 µm ParC layer, as an insulation. This structure matches that used for the NEP. Similar steps and metal thicknesses were used to prepare the TPU-based samples. For this, TPU sheets were laminated on similar glass slides, followed by the deposition of the subsequent metal layers. To achieve TPU thicknesses of more than 25 µm, several 25 µm TPU sheets were laminated together. All samples were subsequently immersed in deionised water to facilitate their release from the substrate. Once fully detached, samples were carefully picked up from the water surface to avoid the formation of wrinkles or mechanical stress. Each sample was then mounted on a custom-designed 3D-printed holder to ensure consistent positioning and tension during acoustic measurements. Material thicknesses were measured using a 3D surface measurement system combining confocal microscopy and interferometry (DCM8, Leica Microsystems, Germany) to measure step thickness and ensure accurate characterisation of each individual layer.

### Ultrasound flow phantom preperation

To estimate the influence of the neural interface on the detection limit of fUS imaging, we prepared a flow phantom model. The phantom was manufactured by pouring 10 wt% gelatin (G9382,Sigma-Aldrich, USA) around a silicone tube (200 µm inner diameter, 200 µm wall thickness) which was connected to openings on opposite sides of a 3D-printed form. The tube was positioned 6 mm beneath the phantom surface. To prepare the gelatin solution, deionised water was heated to 40 °C, and the gelatin powder was gradually added to a large beaker. The mixture was stirred using a magnetic stirrer until fully homogenised, then poured into the mould and left to cool overnight at room temperature. The completed phantom was placed in a water tank, with the tube connected via additional tubing to a flow-inducing pump (36 rpm/0.3 ml*/*min). Blood-mimicking fluid (BMF; 769DF, CIRS, USA) was later pumped through the developed system.

### Flow phantom imaging sequence

Ultrasound images were acquired using an 18 MHz linear array (L22-14vX LF with 128 piezoelectric elements, 100 µm pitch, 22 mm elevation focus, Verasonics, USA) connected to a programmable ultrasound scanner (Vantage 256 High frequency configuration, Verasonics, USA). We transmitted a sequence of 12 angled plane waves (-10° to 10°), resulting in a 1 kHz frame rate after compounding. Ensembles of 200 compounded frames were acquired every 2 s for a total period of 360 s, resulting in an effective power Doppler rate of 0.5 Hz. During acquisition, RF data were real-time delay-and-sum beamformed to complex In-phase and Quadrature (IQ) frames, which were saved for later offline Doppler processing. Real time beamforming was performed using a GPU-accelerated DAS beamformer implemented in CUDA (Quadro P2200, NVIDIA, USA). After acquisition, sets of IQ frames were spatiotemporally filtered using singular value decomposition (SVD) to separate flow signals from the static phantom tissue^70^, after which power Doppler images of the phantom were generated using MATLAB.

### Ultrasound imaging of the flow phantom

In the phantom study, an ultrasound imaging plane was selected to capture the transverse cross-section of the BMF channel. The imaging probe was positioned at three different distances (6,8 and 10 mm) above the channel, and data were recorded in separate sessions for each distance. During each session, the probe was mounted on a motorised XYZ-stage and positioned over the phantom, which was immersed in a water tank. For each session, recordings were made both without and with the NEP placed between the probe and the phantom. To prevent floating, the NEP was weighted down during placement. Additionally, analogous measurements were performed using the patterned electrode array, at an imaging depth of 10 mm to assess the effect of the full device on the reconstructed Doppler signal, while the insulated tracks were placed within the field of view.

### Flow phantom data processing and analysis

We compared recordings obtained without the NEP to those with the NEP placed on top of the phantom. For each scenario, the ultrasound imaging probe was positioned at 6, 8, and 10 mm above the BMF channel to capture a cross-section image of the channel (Fig. 5c). We calculated the mean normalised power Doppler image (MPDI) by averaging across all 180 images. To evaluate the performance of the two different scenarios, we compared the power Doppler intensity, SNR and spatial resolution. We manually identified an ROI by identifying a central point within the channel and included all pixels where the signal amplitude decreased to a threshold of −3 dB. We calculated the normalised power Doppler intensity in decibels of 180 power Doppler images for each scenario (Fig. 5d). To quantify the effect of the NEP, a second ROI corresponding to the flow area was defined and used to estimate the SNR (Fig. 5e) of the power Doppler images as:

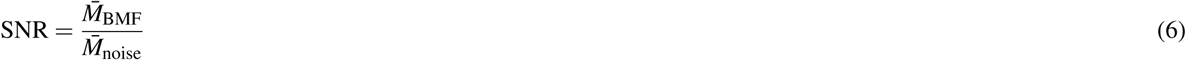

where *M̄* _ROI_ represents the mean power Doppler signal intensity within the flow channel ROI, and *M̄* _noise_ represents the mean signal intensity in the background noise ROI. The flow channel and noise regions are highlighted by the black and white circles, respectively, in the corresponding power Doppler images (Fig. 5c). In addition, resolution is estimated as the FWHM both in axial and lateral directions by assessing the in-plane PSF of the BMF channel. A Gaussian function is fitted to the data along the axial and lateral axes through the centre of the ROI, and the FWHM is calculated. Statistical significance was assessed using the two-sided Student’s t-test. All values were normalised to the mean value of the control (MPDI, with the imaging probe placed 6 mm above the channel and no NEP placed).

### Electrical recordings under imaging conditions

The experiment was conducted similarly to the ultrasound flow phantom study. An ultrasound transparent parylene-based neural electrode array was placed on top of the phantom model and connected to an oscilloscope (RTA4004, Rohde & Schwarz, Germany). Electrode fabrication is detailed in reference^69^. The imaging probe was positioned above the electrode array. Recordings were conducted for four scenarios (Supplementary Fig. 8). (i) The electrode array was placed on the phantom model below the imaging probe within its imaging window. (ii) The electrode array was placed on the phantom model outside of the imaging window. (iii) The electrode array was placed on the phantom model below the imaging probe within the imaging window, and an additional NEP acting as a shield was placed between the imaging probe and electrode array. The additional NEP was connected to the ground of the oscilloscope. (iv) The electrode array was placed on the phantom model below the imaging probe within the imaging window, and the shield-NEP was placed between the imaging probe and electrode array but left floating.

### Experimental model and subject details

The two subject mice were female C57BL/6JRj mice ordered from Jackson Laboratories Netherlands at 7-10 weeks of age, weighing around 20 grams. Mice were socially housed in a 12h:12h light/dark cycle with food and water ad libitum. The in vivo experiments were performed during the dark cycle. All experimental procedures were approved by the Centrale Commissie Dierproeven of the Netherlands (AVD 80110 2020 9725) and by the welfare body of the Netherlands Institute for Neuroscience (IVD, protocol number NIN243601). Studies were conducted in strict accordance with the European Community’s Council Directive (86/609/EEC).

### Surgical procedure

Mice were anesthetized with isoflurane (4-5 % for induction and ∼ 2 % for maintenance) in oxygen (0.8 L*/*min flow rate) though a nose cone. After induction, mice were administered with Buprenorphine (0.1 mg*/*kg, s.c.) and Rimadyl (5 mg*/*kg, s.c.) for analgesia and Dexamethasone (8 mg*/*kg, s.c.) to prevent cerebral oedema and inflammation. During the surgery, lidocaine was applied to the periosteum for local anaesthesia. A cranial window was made over the superior colliculus (SC) by removing the skull while keeping the dura intact. In animal 1, the cranial window extended from 1 mm, to −3.5 anterior-posterior, −1 mm to 3 mm (anterior) and 4 mm (posterior) medial-lateral relative to Bregma a right-angled trapezoid; in animal 2, the cranial window extended from 1 mm to −3.5 mm anterior-posterior, −1 mm to 2.5 (anterior) and 3.5 (posterior) medial-lateral relative to Bregma a right-angled trapezoid. Then, the dura was covered with artificial dura (Cambridge NeuroTech, UK) and sealed by a piece of polymethylpentene (TPX, 250 µm thick, Gooffellow, Germany). A custom-designed head plate was attached to the skull using dental cement (Super-Bond Universal Kit, Sun Medical, Japan). 4-6 h after the surgery, mice were again administrated Buprenorphine (0.1 mg*/*kg, s.c.). They were transferred to a heated recovery chamber, and upon regaining spontaneous mobility, returned to their home cage. 24 h after the surgery, Rimadyl (5 mg*/*kg , s.c.) was administered as post-surgical analgesia.

### Functional ultrasound imaging sequence of in vivo experiments

Similar to the phantom experiments, ultrasound images were acquired using a 18 MHz linear array (L22-14vX with 128 piezoelectric elements, 100 µm pitch, 8 mm elevation focus, Verasonics, USA) connected to a programmable ultrasound scanner (Vantage 256 High frequency configuration, Verasonics, USA). We transmitted a sequence of 12 angled plane waves (-10° to 10°), resulting in a 1 kHz framerate after compounding. Ensembles of 200 compounded frames were acquired every 0.2 s for a total period of 1150 s, resulting in an effective power Doppler rate of 5 Hz. During acquisition, RF data was real time delay-and-sum beamformed to complex IQ frames, which were saved for later offline Doppler processing. Real time beamforming was done using a GPU accelerated DAS beamformer implemented in CUDA (GeForce RTX 3090, NVIDIA, USA). After the recording, IQ frames were spatiotemporally filtered using a singular value decomposition (SVD) to separate blood signals from static tissue, after which the power Doppler image of the phantom was formed using MATLAB.

### Functional ultrasound acquisition of in vivo experiments

Mice were habituated to head-fixation on a running wheel in a dark environment (0-2 lux) by progressively increasing the restraining period from 10 up to 90 minutes over 7 days. Following the habituation period, an experimental session consisting of two recordings was conducted on two consecutive days when the animal was 13 weeks old. Mice were facing a monitor (U2410, Dell, USA) ∼20 cm in front of them. The ultrasound probe is fixed to a micromanipulator, enabling adjustments to target the superior colliculus (SC). Acoustic gel (Aquasonic MAT-00-28152, Parker Laboratories, USA) was applied on the cranial window for ultrasound coupling before placing the ultrasound probe 2 mm above the TPX. For the first recording, functional scans were acquired without the NEP placed on the cranial window. During the second recording, the NEP was placed on the cranial window. The next experiment session was done in a reverse sequence. Each recording consisted of a 10-minute baseline followed by 10 visual stimuli (15 s each), interleaved with 30-second grey screen interstimulus intervals. The visual stimuli consisted of black and white full-field drifting gratings (50 degrees per second, 30-32 lux) moving in one of 8 different directions (angle: 0, 45, 90, 135, 180, 225, 225, 270, 315; randomised sequence) for 1 s each.

### In vivo data processing and analysis

We calculated the mean normalised power Doppler image by averaging across all 5752 images (MPDI). Additionally, we analysed which voxels were modulated by the visual task using a GLM. First, the Doppler data were high-pass filtered (< 0.003 Hz, followed by spatial smoothing (2D Gaussian kernel with *σ* = 1 (FWHM = 2.355 * *σ* )). To generate the GLM regressor for the visual task, the block design was convolved with a canonical hemodynamic response function (HRF), defined by a response onset at 3 s, an undershoot 12 s, a dispersion of 1 s, and a response-to-undershoot ratio of 6. The wheel speed parameters were included in the model as regressors of no interest to reduce potential confounding effects of motion. Effect size was quantified using eta-squared (*η*^2^), which represents the proportion of variance explained by the visual stimulus regressor in GLM. Threshold values of *η*^2^ > 0.1 are interpreted as a significant effect size. Individual effect size images for the control (w/o) and experimental condition (w/) were generated and threshold values were overlaid with the Doppler image (Fig. 6b (i) and Fig. 6c (i)). The effect of the NEP was quantified by measuring the normalised power Doppler intensity within the ROIs across 5752 images acquired under each experimental condition. Similar to the phantom model experiment, we manually selected two areas 300 x 300 mm^2^ within the image (black squares, Fig. 6b(i) and Fig. 6c(i)), a SCR and DBR and included all pixels within the ROI. To complement this, again, we selected two respective regions of interest in the background noise area to estimate the SNR as defined before. We also selected two additional ROIs to calculate the tSNR (Fig. 6f). One in the corresponding area with a high effect size (green squares, Fig. 6b (ii) and Fig. 6c (ii) and one in deeper brain regions with a low effect size (blue squares, Fig. 6b (ii) and Fig. 6c (ii)). For each voxel within the ROI, the time-resolved signal was extracted to obtain voxel-wise time series data. These time series were then averaged across voxels to compute a single representative mean time signal for the ROI. We applied a temporal filter with a moving average (fifteen time points) to the extracted signal. The percentage of change of power Doppler signal in the ROI (Fig. 6b (iii) and Fig. 6c (iii)) is plotted over time for both conditions. tSNR was then calculated as:

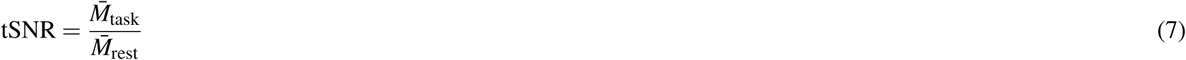

where *M̄* _task_ and *M̄* _rest_ are the means of the Doppler signal during the visual stimulation and resting state, respectively. All values were normalised against the control case (MPDI with no NEP, SCR or ROI I). To assess spatial similarity between Doppler recordings under the two different conditions, we used the SSIM. The SSIM index was then calculated voxel-wise between the MPDI of both conditions to generate a spatial SSIM map. The SSIM is defined as:

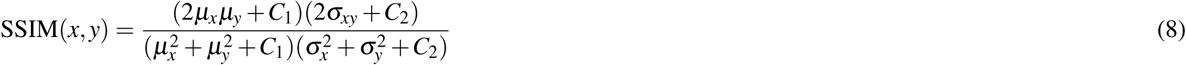

Where *µ_x_*, *µ_y_* are the local means, 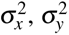 are the local variances and *σ_xy_* is the local covariance between image patches *x*,*y*, obtained from the power Doppler images. *C*_1_ and *C*_2_ are regularisation constants for luminance and contrast. Additionally, to quantify dynamic similarity, each individual frame of the session with the NEP was compared to the control (MPDI with no NEP placed) using the global SSIM index. The global SSIM index between two images is computed as the average of local SSIM values over all *N* pixels.

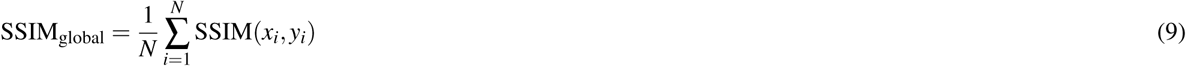

These values were aggregated and presented as a histogram to illustrate the distribution of structural similarity across time. The data were processed using MATLAB.

## Data Availability

All data supporting the findings of this study are available within the article and its Supplementary Information files. Raw datasets generated and/or analysed during the current study have been deposited in the 4TU.ResearchData repository and are publicly available at doi:10.4121/b398b7bc-7f88-4d54-93b0-1ef0b8372392.v2.71

## Code availability

The MATLAB code on which the findings of this study are based is available upon reasonable request to the corresponding author.

## Acknowledgements

This project was financially supported by the Dutch Brain Interface Initiative (DBI2) with project number 024.005.022 of the research programme Gravitation, which is financed by the Dutch Ministry of Education, Culture and Science (OCW) via the Dutch Research Council (NWO) and the Medical Delta Program: Ultrafast Ultrasound for the Heart and Brain program. We thank J. Wilson and S. Tornedde (TBE, Fraunhofer IZM) for providing material samples and for supporting the assembly of the measurement setup. The authors would also like to thank J. Wekselblatt (Ophthalmology, David Geffen School of Medicine, UCLA) for providing the headplate design, and P. Kruizinga (Department of Neuroscience, Erasmus MC) for providing the experimental in vivo fUSI setup (Verasonics system + probe). The authors especially thank M.J. Donahue (Laboratory of Organic Electronics, Linköping University) and D. Byun (Laboratory of Organic Electronics, Linköping University) for providing the parylene-based electrodes.

## Author contributions statement

R.P. drafted the manuscript. R.P. conceptualized the study. R.P. designed the Doppler phantom model. R.P., A.V. and L.H. conducted in vitro experiments. R.W. developed the fUSI sequence. Q.L. conducted the surgery. F.N., Q.L. acquired ethical approval for in vivo experiments C.Q, F.N., Q.L. designed the experimental framework of in vivo experiments. R.P., C.P., Q.L. and C.Q. performed in vivo experiments. F.N. supervised in vivo experiments, C.Q. implemented the GLM and oversaw functional ultrasound imaging and analysis. R.P. processed and analysed the data. V.Gi. and V.Ga. supervised the research. The manuscript was reviewed and edited by R.P., L.H., A.V., C.P., Q.L., F.N., C.Q., R.W., D.M., V.Ga. and V.Gi.

