## Supplementary for "Ultrasound Transparent Neural Interfaces for Multimodal Interaction"

#### ABSTRACT

Neural interfaces that unify diagnostic and therapeutic functionalities hold particular promise for advancing both fundamental neuroscience and clinical neurotechnology. Functional ultrasound imaging (fUSI) has recently emerged as a powerful modality for high-resolution, non-invasive monitoring of brain function and structure. However, conventional metal-based microelectrodes typically impede ultrasound propagation, limiting compatibility with fUSI. Here, we present flexible, ultrasound-transparent neural interfaces that retain practical metal thicknesses while achieving high acoustic transparency. We introduce a theoretical and simulation-based framework to investigate the conditions under which commonly used polymers and metals in neural interfaces can become acoustically transparent. Based on these insights, we propose design guidelines that maximize ultrasound transmission through soft neural interfaces. We experimentally validate our approach through immersion experiments and by demonstrating the acoustic transparency of a suitably engineered interface using fUSI in phantom and in vivo experiments. Finally, we discuss the potential extension of this approach to therapeutic focused ultrasound (FUS). This work establishes a foundation for the development of multimodal neural interfaces with enhanced diagnostic and therapeutic capabilities, enabling both scientific discovery and translational impact.

#### Data Availability

#### Code availability

The MATLAB code on which the findings of this study are based is available upon reasonable request to the corresponding author.

#### Acknowledgements

This project was financially supported by the Dutch Brain Interface Initiative (DBI2) with project number 024.005.022 of the research programme Gravitation, which is financed by the Dutch Ministry of Education, Culture and Science (OCW) via the Dutch Research Council (NWO) and the Medical Delta Program: Ultrafast Ultrasound for the Heart and Brain program. We thank J. Wilson and S. Tornedde (TBE, Fraunhofer IZM) for providing material samples and for supporting the assembly of the measurement setup. The authors would also like to thank J. Wekselblatt (Ophthalmology, David Geffen School of Medicine, UCLA) for providing the headplate design, and P. Kruizinga (Department of Neuroscience, Erasmus MC) for providing the experimental in vivo fUSI setup (Verasonics system + probe). The authors especially thank M.J. Donahue (Laboratory of Organic Electronics, Linköping University) and D. Byun (Laboratory of Organic Electronics, Linköping University) for providing the parylene-based electrodes.

#### Author contributions statement

R.P. drafted the manuscript. R.P. conceptualized the study. R.P. designed the Doppler phantom model. R.P., A.V. and L.H. conducted in vitro experiments. R.W. developed the fUSI sequence. Q.L. conducted the surgery. F.N., Q.L. acquired ethical approval for in vivo experiments C.Q, F.N., Q.L. designed the experimental framework of in vivo experiments. R.P., C.P., Q.L. and C.Q. performed in vivo experiments. F.N. supervised in vivo experiments, C.Q. implemented the GLM and oversaw functional ultrasound imaging and analysis. R.P. processed and analysed the data. V.Gi. and V.Ga. supervised the research. The manuscript was reviewed and edited by R.P., L.H., A.V., C.P., Q.L., F.N., C.Q., R.W., D.M., V.Ga. and V.Gi.

#### Competing Interests

The authors declare no competing interests.

### Supplementary Information

#### Ultrasound Transparent Neural Interfaces for Multimodal Interaction

Raphael Panskus[1,2,*], Andrada I. Velea[1,2], Lukas Holzapfel[2], Christos Pavlou[1], Qingying Li[3], Chaoyi Qin[3], Flora M. Nelissen[3], Rick Waasdorp[4], David Maresca[4], Valeria Gazzola[3,5], and Vasiliki Giagka[1,2,*]

[1]Dept. of Microelectronics, Faculty of Electrical Engineering, Mathematics and Computer Science, Delft University of Technology, Delft, The Netherlands.
[2]Dept. of System Integration and Interconnection Technologies, Fraunhofer Institute for Reliability and Microintegration IZM, Berlin, Germany.
[3]Social Brain Lab, Netherlands Institute for Neuroscience, Royal Netherlands Academy of Art and Sciences, Amsterdam, The Netherlands.
[4]Dept. of Imaging Physics, Faculty of Applied Science, Delft University of Technology, Delft, The Netherlands.
[5]Department of Psychology, University of Amsterdam, Amsterdam, The Netherlands.

[*]

##### Supplementary Note 1

$$|T| = \left[\frac{Z_{\text{in}}}{2}\left(\mathrm{M}_{21} + \frac{\mathrm{M}_{22}}{Z_{\text{out}}} + \frac{\mathrm{M}_{11}}{Z_{\text{in}}} + \frac{\mathrm{M}_{12}}{Z_{\text{in}}Z_{\text{out}}}\right)\right]^{-1} \tag{1}$$

where $\mathrm{M}_{11}$, $\mathrm{M}_{12}$, $\mathrm{M}_{21}$, $\mathrm{M}_{22}$ represent the entities of the total transfer matrix M for $n$ layers:

$$\mathrm{M} = \prod_{i=1}^{n} \begin{bmatrix} \cos(k_i t_i) & jZ_i \sin(k_i t_i) \\ \frac{j\sin(k_i t_i)}{Z_i} & \cos(k_i t_i) \end{bmatrix} = \begin{bmatrix} M_{11} & M_{12} \\ M_{21} & M_{22} \end{bmatrix} \tag{2}$$

The entries for a one-layer system are as follows:

$$M_{11} = \cos(k_1 l_1), \quad M_{12} = jZ_1 \sin(k_1 l_1), \quad M_{21} = \frac{j\sin(k_1 l_1)}{Z_1}, \quad M_{22} = \cos(k_1 l_1)$$

For multilayer systems let's define:

$$c_i = \cos(k_i t_i)$$
$$s_i = \sin(k_i t_i)$$

Therefore, the matrix entities for a three-layer system can be expressed as:

$$M_{11} = c_1c_2c_3 - \frac{Z_1}{Z_2}s_1s_2c_3 - \frac{1}{Z_3}(Z_2c_1s_2 + Z_1s_1c_2)s_3$$

$$M_{12} = j\left[Z_3c_1c_2s_3 - Z_3\frac{Z_1}{Z_2}s_1s_2s_3 + Z_2c_1s_2c_3 + Z_1s_1c_2c_3\right]$$

$$M_{21} = j\left[\left(\frac{s_1c_2}{Z_1} + \frac{c_1s_2}{Z_2}\right)c_3 + \frac{1}{Z_3}\left(c_1c_2s_3 - \frac{Z_2}{Z_1}s_1s_2s_3\right)\right]$$

$$M_{22} = jZ_3s_3\left(\frac{s_1c_2}{Z_1} + \frac{c_1s_2}{Z_2}\right) + c_1c_2c_3 - \frac{Z_2}{Z_1}s_1s_2c_3$$

The final matrix entities of the four-layer system are as follows:

$$M_{11} = \frac{-Z_3\left(Z_2\left(Z_1 s_1 c_2 + Z_2 s_2 c_1\right) c_3 - Z_3\left(Z_1 s_1 s_2 - Z_2 c_1 c_2\right) s_3\right) s_4 + Z_4\left(Z_2\left(Z_1 s_1 c_2 + Z_2 s_2 c_1\right) s_3 + Z_3\left(Z_1 s_1 s_2 - Z_2 c_1 c_2\right) c_3\right) c_4}{Z_2 Z_3 Z_4}$$

$$M_{12} = \frac{i\left[Z_3\left(Z_2\left(Z_1 s_1 c_2 + Z_2 s_2 c_1\right) c_3 - Z_3\left(Z_1 s_1 s_2 - Z_2 c_1 c_2\right) s_3\right) c_4 - Z_4\left(Z_2\left(Z_1 s_1 c_2 + Z_2 s_2 c_1\right) s_3 + Z_3\left(Z_1 s_1 s_2 - Z_2 c_1 c_2\right) c_3\right) s_4\right]}{Z_2 Z_3}$$

$$M_{21} = \frac{i\left[Z_3\left(Z_2\left(Z_1 c_1 c_2 - Z_2 s_1 s_2\right) c_3 - Z_3\left(Z_1 s_2 c_1 + Z_2 s_1 c_2\right) s_3\right) s_4 + Z_4\left(Z_2\left(Z_1 c_1 c_2 - Z_2 s_1 s_2\right) s_3 + Z_3\left(Z_1 s_2 c_1 + Z_2 s_1 c_2\right) c_3\right) c_4\right]}{Z_1 Z_2 Z_3 Z_4}$$

$$M_{22} = \frac{Z_3\left(Z_2\left(Z_1 c_1 c_2 - Z_2 s_1 s_2\right) c_3 - Z_3\left(Z_1 s_2 c_1 + Z_2 s_1 c_2\right) s_3\right) c_4 - Z_4\left(Z_2\left(Z_1 c_1 c_2 - Z_2 s_1 s_2\right) s_3 + Z_3\left(Z_1 s_2 c_1 + Z_2 s_1 c_2\right) c_3\right) s_4}{Z_1 Z_2 Z_3}$$

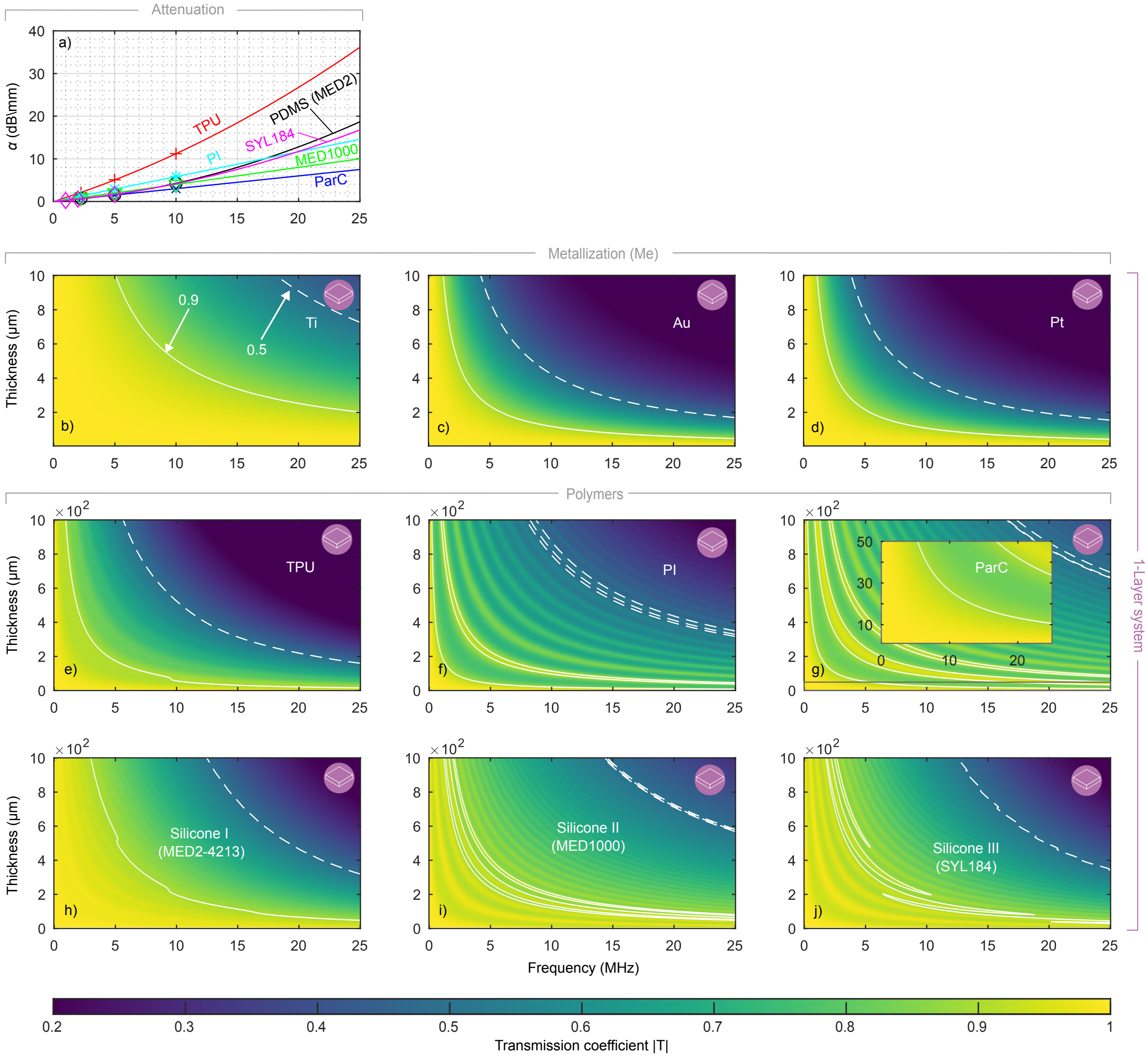


**Supplementary Figure 1. Transmission maps of polymers and metals.** a) Frequency-dependent acoustic attenuation coefficients for selected polymers compiled from literature. Solid lines denote quadratic fits to the reported data. b–j), Numerical simulations of the ultrasound transmission coefficient as a function of frequency and thickness for single-layer systems of Ti (b), Au (c), Pt (d) TPU (e), PI (f), ParC (g), Silicone I (MED2-4213 (h), Silicone II (MED-1000) (i) and Silicone III (SYL184) (j). Solid contours indicate 0.9 transmission, and dashed contours indicate 0.5 transmission.

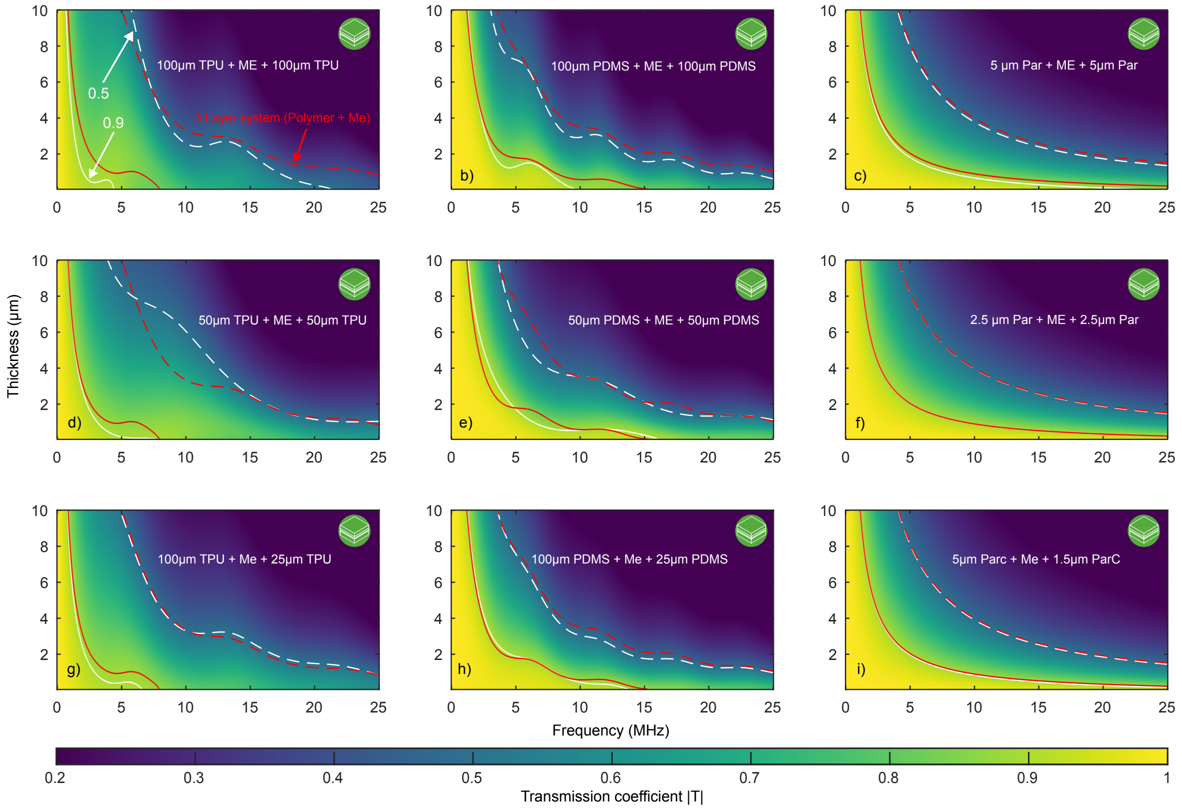


**Supplementary Figure 2. Transmission maps for four-layer systems.** a–c), Simulated transmission maps for four-layer stacks comprising a polymer substrate and insulation of the same thickness (100 μm TPU, PDMS, or 5 μm ParC). d–f), Simulated transmission maps for four-layer stacks comprising a polymer substrate and insulation of the same thickness simultaneously keeping the overall polymer thickness comparable (50 μm TPU, PDMS, or 2.5 μm ParC). g–i) Transmission maps for four-layer systems incorporating a thin insulation layer (100 μm TPU, PDMS, or 5 μm ParC substrate and 25 μm TPU, PDMS, or 1.5 μm ParC insulation). Solid contours indicate 0.9 transmission, and dashed contours indicate 0.5 transmission. Dashed red contours indicate the transmission threshold of the corresponding three-layer configurations, for direct comparison.

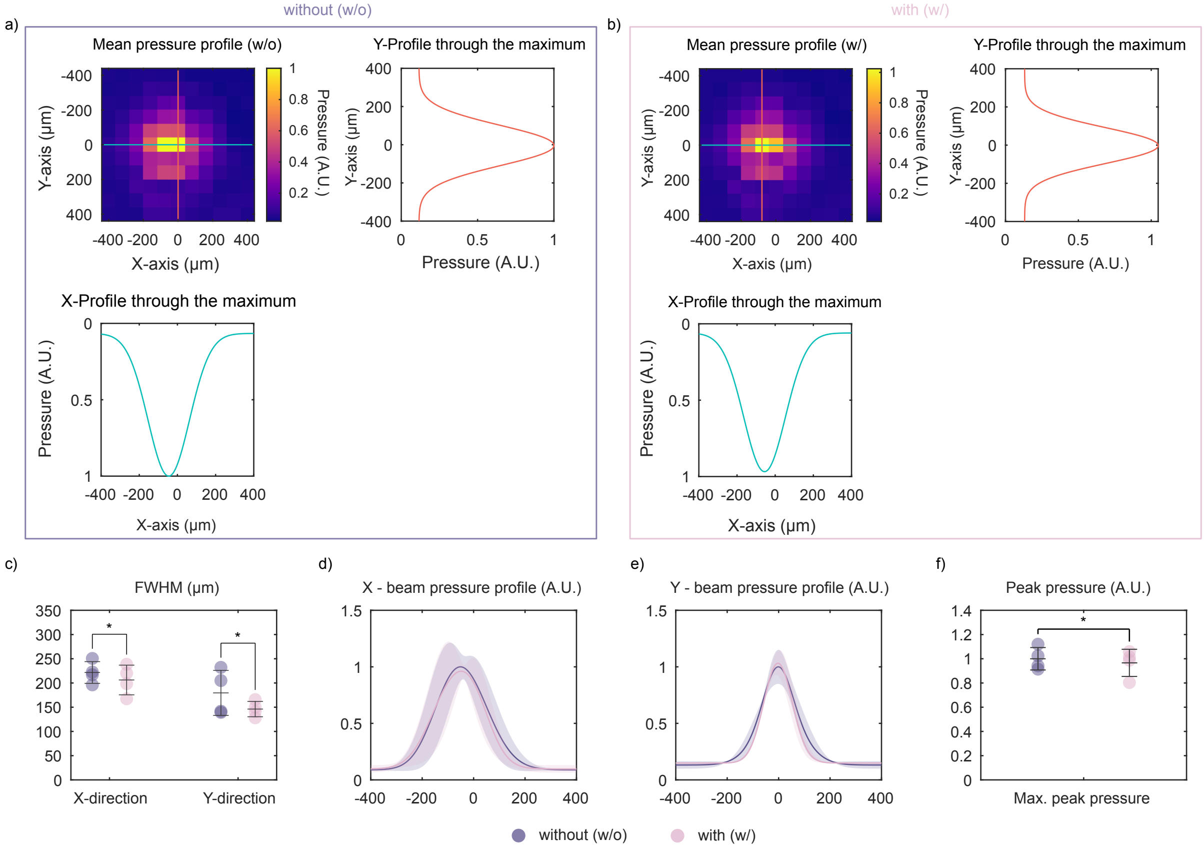


**Supplementary Figure 3.** **Ultrasound beam profile characterisation of the FUS transducer with NEP.** Normalised acoustic pressure distributions in the Y-X section of the focus region and corresponding X-and Y-profiles without (a) and with (b) NEP. FWHM of the beam profiles in X and Y directions (c). Fitted curves to the extracted X- (d) and Y-profiles (e), and normalised peak pressure (f). Error bars and shading represent variability across repeated scans (*P > 0.05).

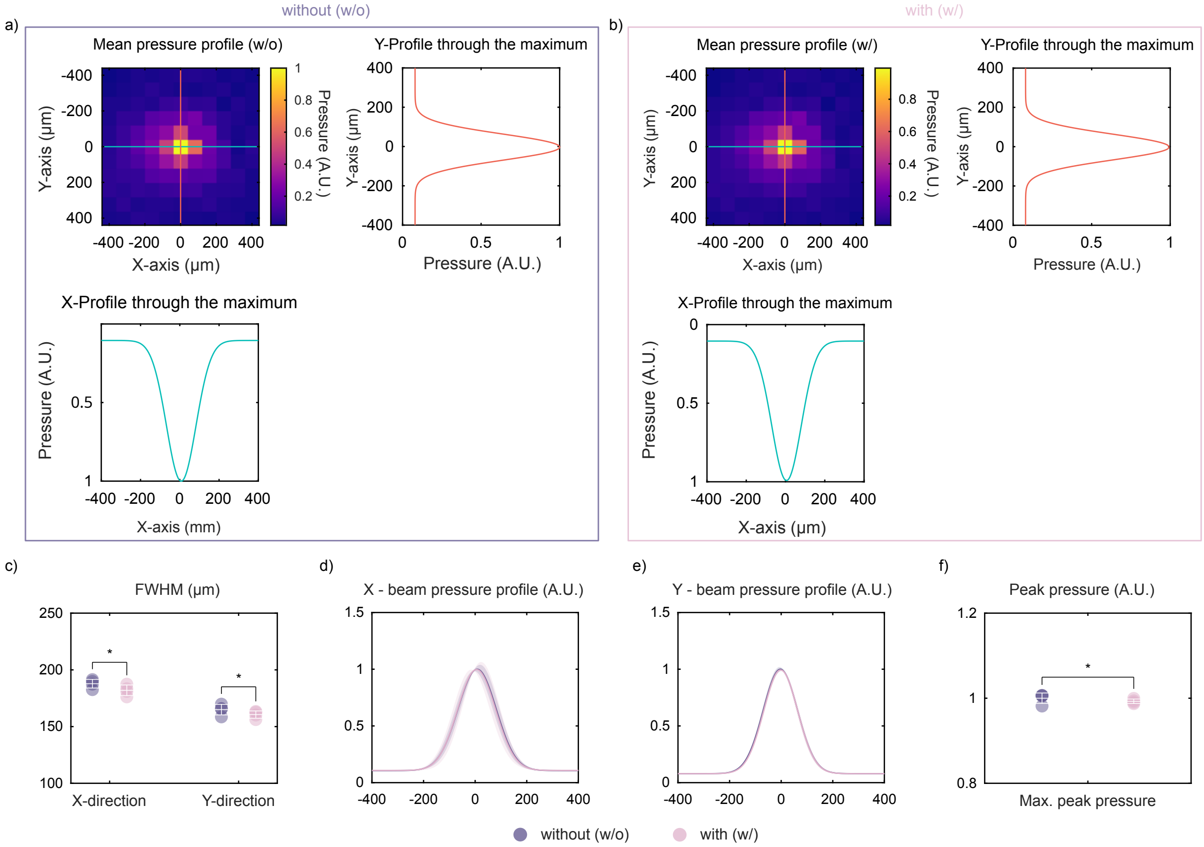


**Supplementary Figure 4. Ultrasound beam profile characterization of the FUS transducer with a parylene based electrode array.** Normalized acoustic pressure distributions in the Y–X section of the focus region and the corresponding X- and Y-profiles without (a) and with (b) the electrode array. FWHM of the beam profiles in the X and Y directions (c). Fitted curves to the extracted X- (d) and Y-profiles (e), and normalized peak pressure (f). Error bars and shading represent variability across repeated scans (*P > 0.05).

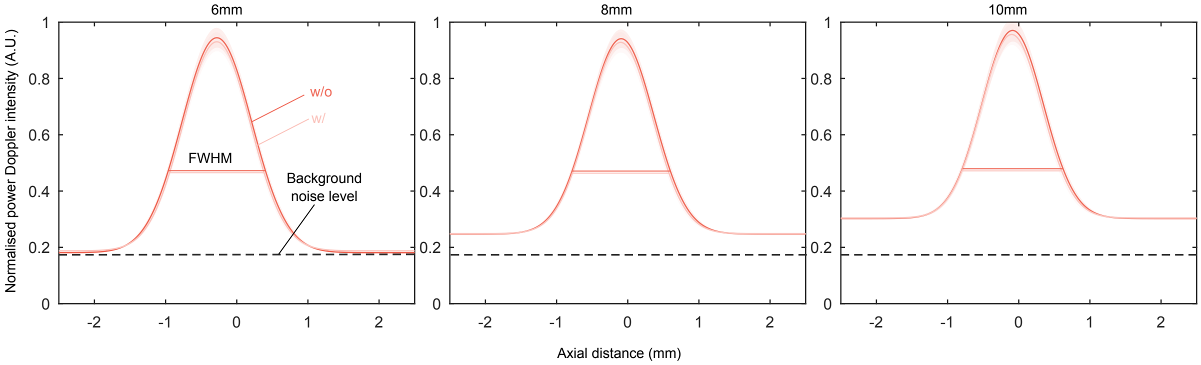


**Supplementary Figure 5. Axial PSF profile.** a-c) represent the fitted curve to the mean axial PSF cross-section of the BMF channel at each imaging depth. (a) axial PSF profile at 6 mm. (b) axial PSF profile at 8 mm. (c) axial PSF profile at 10 mm. Dashed black contours indicate the background noise at a depth of 6 mm. Shaded areas represent the standard deviation. Power Doppler data are normalised against the mean of the control (no NEP, 6 mm).

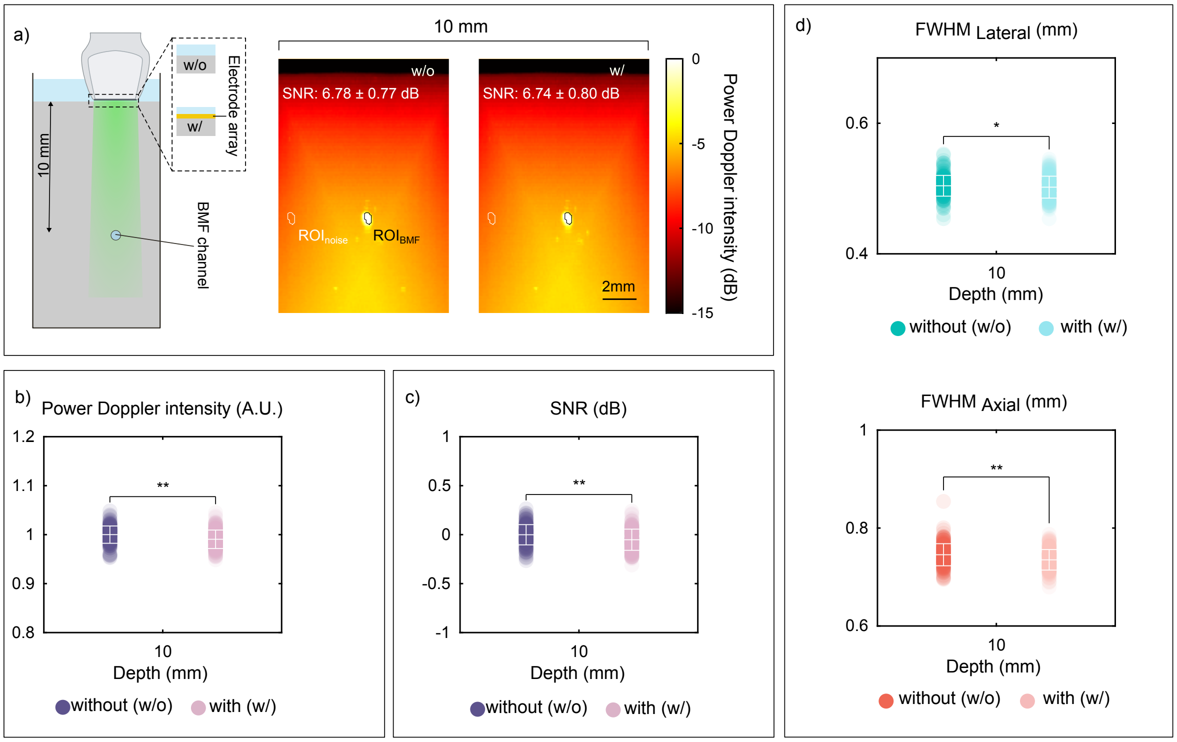


**Supplementary Figure 6. Compatibility of functional ultrasound imaging with a parylene based electrode array using a flow phantom model.** (a) Schematic of the data recording principle with the US probe placed at an imaging depth of 10 mm above the BMF and MPDI of the blood flow phantom, acquired without (w/o) and with (w/) the electrode array. The SNR within the ROI is reported as mean ± standard deviation (SD). Power Doppler intensity (b) and SNR (c) of the ROI with and without the electrode array. (d) Characterization of lateral and axial resolution. Data are presented as mean ± SD; dots represent individual acquisitions, n = 180 acquisitions per group. All values were normalized against the control (MPDI, no electrode array). $^{*}P > 0.05$, $^{**}P < 0.01$.

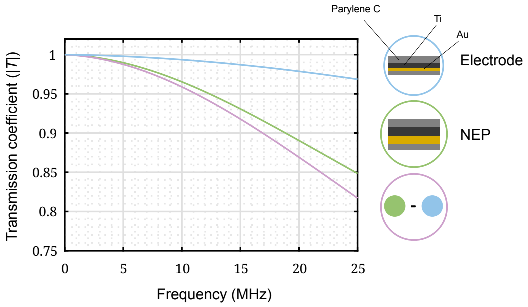


**Supplementary Figure 7. Simulated transmission coefficients plotted over frequency.** Simulated transmission coefficients for the electrode array (blue), the NEP (green), and their combined configuration (purple).

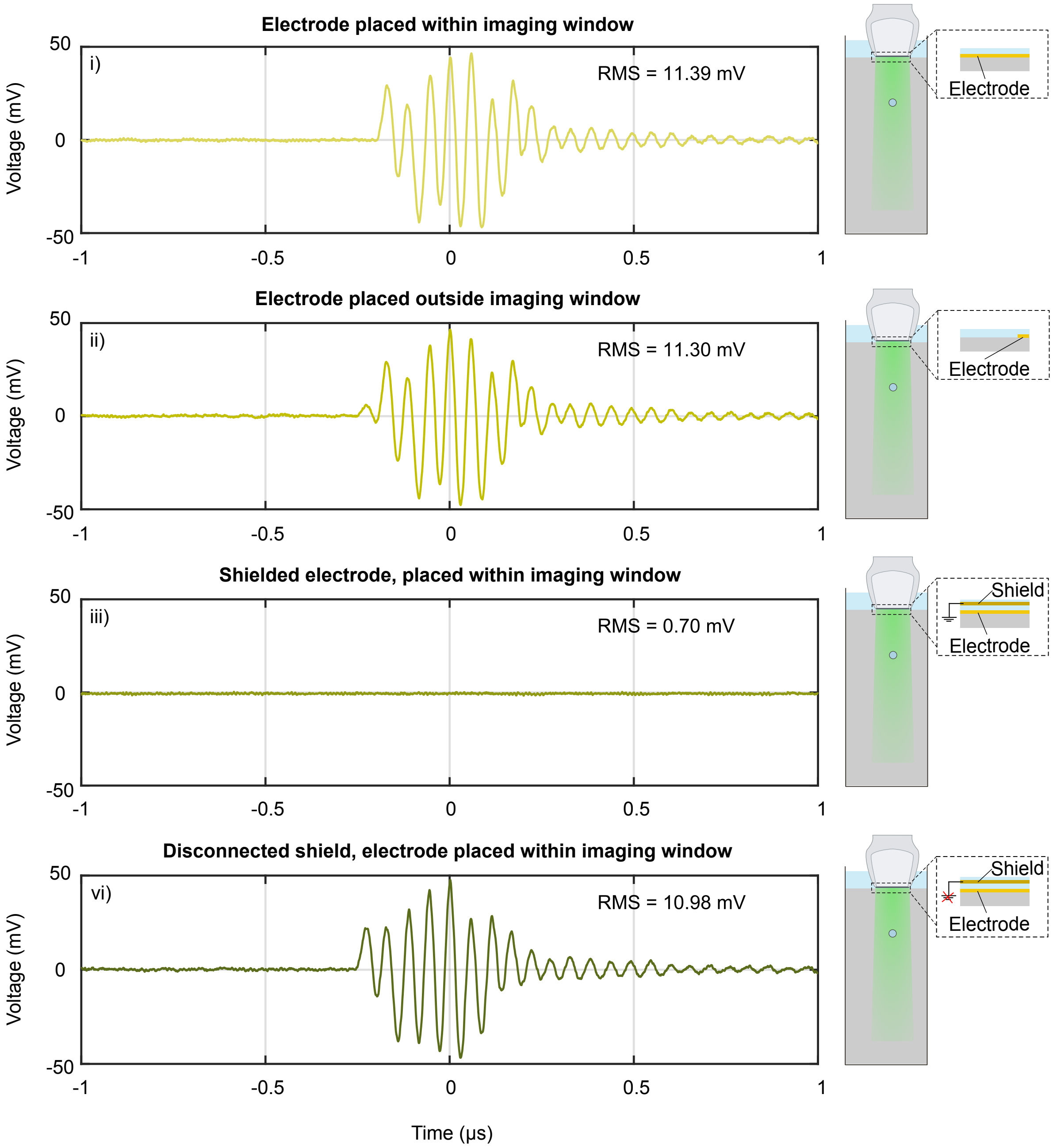


**Supplementary Figure 8. Recordings with a parylene-based electrode during fUSI of a phantom.** Recordings were obtained with the electrode connected to an oscilloscope under four conditions: (i) electrode array placed within the imaging window; (ii) electrode array outside the imaging window; (iii) electrode array within the imaging window with a ground-connected shield; and (iv) electrode within the imaging window with a floating shield. Scenarios (i), (ii) and (iv) show the captured ultrasound pulse of the imaging probe due to electrical coupling. Notably, no pulse was detected in scenario (iii), where the electrode was positioned within the imaging window and appropriately shielded. Careful shielding can avoid artifacts on the electrical recording due to coupling from the imaging system (iii).

#### Supplementary Note 2

For neural interfaces, materials are chosen based on the specific requirements of flexibility, biocompatibility, mechanical robustness and electrical performance for long-term neural interfacing. Generally, thicknesses are dependent on the desired mechanical properties and deposition method.

- Parylene C is commonly used for substrate and insulating layers due to its excellent biocompatibility and conformal coating capabilities. For neural interfaces, practical thicknesses typically range from 1 to 10 µm[1,2]. This is due to the chemical vapor deposition reaching thickness within the nm-µm range[3]. Due to the mechanical properties, thicker layers lead to stiff and brittle probes.

- PDMS (Polydimethylsiloxane) is widely used in flexible neural interfaces due to its softness and elasticity. The practical thickness for PDMS in neural applications ranges from several µm to hundreds of µm[4–7], depending on the desired flexibility and mechanical properties. Thicker PDMS layers are often used in more robust, soft interfaces, while thinner layers may be applied when flexibility and stretchability are prioritised. Typically, several hundred µm can be achieved by spin-coating PDMS films.

- TPU (Thermoplastic Polyurethane) is a relatively new material compared to PDMS or ParC, and is not yet as widely used in neural interfaces. The thickness of TPU films typically falls within a similar range to that of PDMS, owing to their comparable mechanical properties. This thickness range strikes a balance between mechanical strength, flexibility, and the ability to conform to neural tissue without compromising device performance. In this work, we have used commercially available TPU foils (which typically start at 25 µm in thickness). However, we have also fabricated custom, TPU films via spin-coating, ranging from several nm to several µm.

- PI (Polyimide) is a widely used polymer in neural interface technology due to its excellent biocompatibility, flexibility, and chemical stability. Typically, polyimide is fabricated using spin coating, where a liquid precursor is deposited onto a substrate and then thermally cured to form a solid, imidized film with a possible thickness between one and tens of µm[8].

- Hydrogels are increasingly being explored for neural interfaces due to their unique ability to closely match the mechanical properties of brain and nerve tissue. Their high water content and softness help minimise the mechanical mismatch between implants and surrounding neural tissue, reducing inflammation and improving long-term biocompatibility. In neural applications, hydrogels can be engineered to be electrically conductive or to deliver bioactive molecules, enhancing signal transmission and promoting tissue integration. They are typically fabricated using methods like photopolymerization or chemical crosslinking, with thicknesses ranging from a few micrometers to several hundred micrometers, depending on the application[9].

- Metallization layers are traditionally used to form electrodes and interconnects that transmit electrical signals between the device and neural tissue. Commonly used metals include Au, Pt, and Ti due to their excellent electrical conductivity, corrosion resistance, and biocompatibility. These metal layers are typically deposited via sputtering or evaporation and patterned using photolithography. Thicknesses usually range from 100 nm to 1 µm, balancing electrical performance with flexibility. For applications requiring lower impedance or enhanced charge injection capacity, such as stimulation electrodes, thicker metal layers (up to 5–20 µm) can be achieved through electroplating. However, metals are significantly stiffer than the surrounding polymers or tissue, which can lead to a mechanical mismatch. To mitigate this, metals are often thin, embedded within flexible substrates or patterned in serpentine shapes to enhance stretchability and reduce stress on neural tissue.

- Graphene is gaining increasing attention for neural electrode applications due to its high carrier mobility, mechanical flexibility, and biocompatibility. While its absolute electrical conductivity is lower than that of bulk metals, graphene's performance as a conductive and ultrathin material makes it particularly well-suited for bioelectronic interfaces. Its atomically thin, transparent structure allows for highly sensitive neural recording with minimal tissue disruption. Graphene electrodes can conform closely to neural tissue, reducing inflammation and improving long-term stability. Typically, graphene is deposited using CVD and then transferred onto flexible substrates, forming layer thicknesses ranging from sub-nm to a few nm. These properties position graphene as a next-generation material for developing ultra-flexible, high-performance neural interfaces[10,11].

For ultrasound-transparent electrodes, materials with acoustic impedance closely matched to the surrounding medium and low attenuation coefficients are generally preferred to maximise acoustic transmission. However, limitations in these properties can be partially mitigated by minimising material thickness, provided that the material's Young's modulus allows for mechanical stability and low flexural rigidity. Ultimately, the dominant factors influencing acoustic transparency include the acoustic impedance mismatch, attenuation, and the material's ability to form thin, mechanically stable layers. Balancing these parameters is key to optimising both acoustic and mechanical performance in neural interface applications. Supplementary Tab. 1lists the key material properties with respect to the acoustic and mechanical performance of neural electrodes.

| Materials | | Attenuation | Acoustic Impedance | Young's Modulus | Typical thicknesses (µm) |
|---|---|---|---|---|---|
| Polymers | TPU | - | + | + | 1–100 (spin coating) |
| | Silicone | + | ++ | ++ | 1–100 (spin coating) |
| | Parylene | ++ | - | - | 1–50 |
| | Polyimide | - | - | - | 1–15 |
| Metals | | +++ | - - | - - | 0.05–15 |
| PEDOT:PSS | | + | + | - | 0.1–1 |
| Graphene | | - - | - - | - - | 0.001–0.1 |
| Hydrogels | | + | ++ | +++ | 10–1000 |

**Supplementary Table 1.** Qualitative comparison of materials and their material properties with respect to the acoustic and mechanical performance of neural electrodes.
